# Deciphering the functional association of novel variants of *BMP7* in isolated congenital heart disease by integrating *in vitro* and *in silico* approaches

**DOI:** 10.64898/2025.12.23.696272

**Authors:** Jyoti Maddhesiya, Ritu Dixit, Ashok Kumar, Bhagyalaxmi Mohapatra

## Abstract

Bone morphogenetic protein7 (BMP7), an important member of the TGFβ superfamily, is known to be vital for embryonic growth and development. To decipher the role of BMP7 in congenital cardiac malformations, genetic screening of 285 CHD cases along with 400 healthy controls individuals was performed by Sanger sequencing method. Five missense variants were identified in 5 unrelated CHD probands with distinct phenotypes. Three novel missense (p.D85V, p.R175W, and p.A283T) variants in the pro-peptide region and two other variants (p.M315I and p.N321S) in the mature domain were documented. *In vitro* functional analysis revealed WT as well as all five mutant BMP7 proteins localized within ER compartment, which was confirmed by ER-Tracker and SERCA2 staining. Western blot analysis demonstrated enhanced phosphorylation of SMAD1/5 associated with these variants. Furthermore, transactivation assays showed increased activity of BMP-responsive promoters-*Id1-luc*, *Id3-luc*, *Tlx2-luc*, and *p(SBE)4-luc* with a synergistic effect observed upon *AKL2* co-expression. Additionally, overexpression of downstream targets, namely *Bmp2*, *Bmp7*, *Nkx2.5*, *Gata4*, *Irx4*, *Smad1*, *Smad4*, and *Smad5,* alongside downregulation of left-right patterning genes (*Nodal* and *Pitx2*) and the BMP antagonist *Chordin,* was observed. Likewise, EdU staining revealed increased cellular proliferation associated with all these variants. Moreover, modeling of secondary and tertiary structures also suggested that these variants might induce conformational changes in BMP7, potentially strengthening its binding affinity with the receptor and thereby amplifying SMAD signaling. Collectively, *in vitro* and *in silico* analyses suggest that these *BMP7* variants exhibit gain-of-function (GoF) activity, disrupting normal BMP signaling pathways and thereby contributing to the development of CHD.

## 1. Introduction

Congenital heart diseases (CHDs) are a group of structural and functional defects in the cardio-circulatory system that occur during the development of the heart and affect ∼1% of all live newborns (Thomford *et al*., 2018). Animal models as well as clinical genetic studies provide evidence for the genetic contribution of congenital heart disease. The cardiogenesis process is coordinated through the sequential and precise spatiotemporal control of expression of multiple genes including cardiac-rich transcription factors *NKX2.5* (McElhinney *et al*., 2003), *GATA4* (Butler *et al*., 2010), *TBX20* (Posch *et al*., 2010), signalling molecules namely *TGFβ* (Shi *et al*., 2019), *BMPs* (Wang *et al*., 2011) and *Nodal* (Mohapatra *et al*., 2009) and structural proteins such as *MYH7, MYH6* and *ACTC1* (Maddhesiya and Mohapatra, 2024) those converge into multiple synchronized signalling pathways. However, the genetic determinants in the majority of cases remain unclear, as CHD is a genetically heterogeneous malady.

Bone morphogenetic protein 7 (BMP7), originally discovered for its ability to induce bone formation, also known as osteogenic protein1 (OP1), is a member of TGFβ super-family. This is a dimeric secretory molecule, widely expressed in embryonic tissue; however further studies have shown its importance in other organ system development as well (Hogan, 1996). It is well established that BMP7 forms heterodimers with either BMP2 or BMP4 (Kim *et al*., 2019). These are known to be crucial players in a wide range of biological processes involving cellular differentiation, proliferation, tissue regeneration, and migration of mesenchymal stem cells (MSCs) into chondroblasts (Mukhametov *et al*., 2022; Sampath and Vukicevic, 2020). During cardiogenesis, BMP signalling has a critical role in SHF specification, regulation of proliferation, left-right patterning, and induction of myocardial differentiation (Wang *et al*., 2011). The role of BMP signalling for valve formation is defined in two phases: first it activates the endocardium and later coordinates with TGFβ and Notch signalling to induce endothelial-to-mesenchymal transformation (EMT) followed by incursion of mesenchyme into the cardiac cushion (Garside *et al*., 2013).

The role of BMPs in heart development was first determined in dorsal vessel formation of *Drosophilla Melanogaster*. Several BMPs, like BMP2, 4, 5, 6, 7 and 10, show their expression in the AV cushion and OFT during valve formation, where they function independently and redundantly, which strongly affect the maturation of these tissues in the heart (Yuasa and Fukuda, 2008). These BMPs are strongly expressed in the developing myocardium of 8.5 dpc mouse embryos (Jones *et al*., 1991; Lyons *et al*., 1995; Dudley and Robertson, 1997); *Bmp7*-deficient mice die shortly after birth and display severe defects of kidney and eye development, as well as minor skeletal defects (Dudley and Robertson, 1997; Luo *et al*., 1995). *Bmp5*, *6*, and *7* are grouped together and co-express during cardiogenesis, since individual knockout (KO) of these failed to uncover its role in heart development. However, double KO mice are observed with severe phenotypes. Compound *Bmp5* and *Bmp7* mutant mice develop with defective chamber septation with myocardial and pericardial abnormalities (Solloway and Robertson, 1999). Similarly, delay in the formation of the outflow tract (OFT) endocardial cushions was noted in *Bmp6*;*Bmp7* double mutant mice and these mutant mice die between 10.5 and 15.5dpc. Since only a fraction of these *Bmp6*;*Bmp7* mutant mice show defects in valve morphogenesis and chamber septation(Kim *et al*., 2001), histological analysis of these double mutants embryo revealed defective chamber septation and hypoplastic ventricles with reduced trabeculations (Kim *et al*., 2001). The expression of BMP7 was found to be high in the OFT myocardium of OFT-restricted *Bmp4* (KO) mice; however, *Bmp7* and OFT-restricted BMP4 double (KO) mice display severe reductions in the OFT cushions and more severe shortening of the OFT when compared with the OFT-restricted *Bmp4* single (KO) mice (Yuasa and Fukuda, 2008).

The commencement of BMP signalling occurs after binding of BMP7 ligand with their receptor BMPR2 (ALK2), on the cell surface followed by phosphorylation of regulatory SMAD1, SMAD5 and SMAD8. Phosphorylated SMAD1 and SMAD5 form a complex with SMAD4 to transduce a signal by bonding and activating different downstream target gene promoters such as *Id1* (Kantzer *et al*., 2022; Du *et al*., 2010; Weiskirchen *et al*., 2009; Izumi *et al*., 2006), *Id2*(Miyazono *et al*., 2005; Du *et al*., 2022), *Id3*, *Id4*, *Tlx2*, *Xvent2* and *Runx2*(Mukhametov *et al*., 2022; Aluganti Narasimhulu and Singla, 2020). Additionally, BMP signalling collaborates with the Nodal signalling pathway in order to regulate the differentiation and proliferation of early mesodermal and cardiac progenitor cells by monitoring the expression of cardiac-enriched transcription factors such as *Gata4, Nkx2-5, Irx4*, *Tbx5, Mef2c, Sox4,* and *Tnntt2* in mES cells (Cai *et al*., 2013; Chhabra *et al*., 2019). Besides the SMAD-dependent canonical pathway, BMP7 can trigger SMAD-independent non-canonical signaling routes, such as the MAP kinase pathway through the serine–threonine kinase TAK1 (Goswami *et al*., 2001; Wang *et al*., 2014). It also plays a role in the activation of the PI3K/Akt and Rho-GTPase pathways (Wang *et al*., 2014).

Although increasing evidence suggests an essential role of BMP7 during heart development, the association of BMP7 mutations with cardiac malformation is lacking. Further, the mechanism underlying the *BMP7* mediated CHD phenotype is not clearly understood. Molecular genetic investigations are required to provide precise insights into association of BMP7 with *CHD*. This prompted us to perform a detailed molecular genetic analysis of *BMP7* in a cohort of 285 CHD cases, which resulted in identification of five non-synonymous, variants that were not found in 4 hundred controls from our population, suggesting the possible disease-association of these variants. Further, comprehensive *in vitro* and *in silico* analyses of all missense variations suggested that BMP7 could be a causative factor for CHD.

## 2. Materials and Methods

### 2.1 Clinical evaluation

We have recruited a cohort of 285 patients (174 males and 111 females), which include 8 familial and 277 sporadic cases (with a median age of 3 years), from the Departments of Pediatrics and Cardiovascular & Thoracic surgery, Institute of Medical Sciences, BHU, Varanasi. 400 healthy, age-matched control individuals (median age; 3.7 years) with no history of CHD were also enrolled for this study from the same geographical locality after obtaining the written informed consent. A thorough, comprehensive clinical evaluation, including detailed individual and familial histories, chest radiography, ECG and 2D-colour Doppler echocardiography was performed to confirm the disease condition. Probands with known chromosomal abnormalities or syndromic CHD cases were excluded from the study. The study design was approved by the ‘Institutional Ethical Committee’ of the University. Probands enrolled in the study had a wide spectrum of CHD phenotypes; the clinical phenotypes of the probands along with their minor allele frequency (MAF) are listed in the **(Supplemental Table S1).**

### 2.2 Mutational screening and variant analysis

Genomic DNA was extracted from peripheral blood (3-5 ml) leukocytes from all subjects by standard ethanol precipitation protocol. Primers were designed to amplify the coding exons and splicing boundaries of the human *BMP7* (NG_032771.1) by polymerase chain reaction (PCR). The amplicons were purified by enzymes Exonuclease I (USB Products, Affymetrix, Inc., USA) and recombinant Shrimp Alkaline Phosphatase (USB Products, Affymetrix, Inc., USA) as per the manufacture’s protocol. Purified PCR products were Sanger sequenced using the Big Dye® Terminator v3.1 Cycle Sequencing Kit (Applied Biosystems, Inc., USA) on genetic analyzer by Applied Biosystems (ABI PRISM 3130, USA). The DNA sequences in electropherograms were analyzed by Finch TV software (http://www.geospiza.com/ftvdlinfo.html, Geospiza). Each detected variant was confirmed by re-sequencing of the variation carrying DNA samples with alternate primers as well as by sequencing of independently PCR-amplified products of the same exon from the respective subject. The parental genotyping was also done to check the inheritance pattern of the variant. To confirm the novelty of identified variants, gnomAD (https://gnomad.broadinstitute.org/), ClinVar (http://www.ncbi.nlm.nih.gov/clinvar/), 1000G (https://www.internationalgenome.org/) and dbSNP (https://www.ncbi.nlm.nih.gov/snp/) as well as databases from the same locality INDEX-db (http://indexdb.ncbs.res.in/), GenomeAsia100K (https://browser.genomeasia100k.org/) were queried.

### 2.3 In silico analysis of the variants

Different bioinformatic tools were used for predicting the pathogenicity, stability, structure, and functions of wild-type (WT) and mutant-proteins (muteins) for identified variants. *In-silico* analyses of *BMP7* were performed by retrieving the mRNA sequence (NM_001719.3) and protein sequence (NP_001710.1) from NCBI (https://www.ncbi.nlm.nih.gov/genbank/) and the Protein database (https://www.ncbi.nlm.nih.gov/protein/).

#### 2.3.1 Multiple Sequence Alignment of conserved amino acids

Multiple sequence alignment of BMP7 protein was performed across different orthologues species including *H.sapiens, P.troglodytes, M.mulatta, C.lupus, B.taurus, M.musculus, R.norvegicus, G.gallus, D.rerio* to check the conservation of mutated residues by ‘Homologene’ tool of NCBI (http://www.ncbi.nlm.nih.gov/homologene).

#### 2.3.2 Prediction of potential pathogenicity of identified variants of BMP7

The disease-causing potential as well as structural alteration, stability and function of proteins was predicted by an integrated clinical and genetic database that incorporates more than 15 *in-silico* tools such as SIFT, Polyphen2_HVAR, LRT, MutationTaster, PROVEAN, VEST3, M_CAP, CADD, DANN, GenoCanyon, ClinPred, ReVe, REVEL etc. These bioinformatic tools are based on different parameters and algorithms, on the basis of which these can predict the effect of missense variation on the conservation of nucleotides, amino acid residues, and the nature of R-groups.

#### 2.3.3 Prediction of 2-D and 3-D structures of BMP7 muteins

To predict the structural changes introduced into the muteins secondary structures caused by non-synonymous mutations, the Psipred server (http://bioinf.cs.ucl.ac.uk/psipred/) was used. To further validate these conformational changes, modelling of tertiary structures of BMP7 WT and all muteins was performed by Alphafold2 googlecolab, (https://colab.research.google.com/github/sokrypton/ColabFold/blob/main/AlphaFold2.ipyn). To visualize the differences in the conformation of the modelled WT versus muteins, PyMOL Molecular Graphics System, Version 3.1 (Schrödinger, LLC.) was used to align the two comparable structures. Furthermore, to validate the visualized differences, global RMSD at alpha carbon was calculated by UCSF ChimeraX.

### 2.4 In vitro functional analysis of variants

#### 2.4.1 Cloning of BMP7 into expression vector followed by site-directed mutagenesis

The WT full length MGC clone of human *BMP7* ORF (Acc no. BC008584 BF338314) was purchased from “Thermo Scientific” (USA). The coding region including Kozak sequence was PCR amplified using the Pfu-Ultra high-fidelity DNA polymerase (Agilent, Santa Clara, CA, USA) and subcloned into pcDNAv3.1/V5-His TOPO TA vector (Invitrogen, Carlsbad, CA, USA). The identified variants were introduced into the WT-BMP7-pcDNAv3.1 to prepare the mutant (MUT) constructs (c.254A>T; p.D85V, c.523C>T; p.R175W, c.847G>A; p.A283T, c.945G >A; p.N315I, c.962A>G; p.N321S) by using Quick change II XL site-directed mutagenesis kit (Agilent Technologies Inc., USA) with a complementary pair of primers (as per manufacturer’s descriptions). These MUT constructs were transformed into XL10 gold competent cells followed by plasmid isolation (Qiagen midi kit, Germany). The sequence integrity of all MUT constructs were confirmed by Sanger sequencing.

#### 2.4.2 Cellular culture and DNA transfection

For *in vitro* functional analysis, mouse pluripotent embryonic cell line P19 (kind gift from Dr. Ramkumar Sambasivan, InStem, India) was grown in α-MEM (Gibco, Life technologies Corp., USA) supplemented with 10% fetal bovine serum (FBS, HIMEDIA, India.) with 100 units/mL penicillin and 100 units/mL streptomycin. The H9c2 cell-line (purchased from cell repository, NCCS, India) were cultured in DMEM+10% FBS at 37°C with 5% CO2 in a humid atmosphere. Immunostaining was performed in H9c2 cells as these are cardiomyoblast cells from embryonic rat, and having better morphology than P19 which could help in analysis of any cytoskeletal deformities. Transfection for each experiment was performed after 24 hrs of cells seeding at 30-40% confluency with FuGENE6 transfection reagent (Promega Corp., IN, USA) as per manufacturer’s protocol.

#### 2.4.3 Immunocytochemistry

For visualizing the expression and cellular localization of BMP7, immunocytochemistry of WT and BMP7 muteins was performed in P19 as well as H9c2 cell lines in independent experiments. Both types of cells were grown on coverslips placed in 6-well culture plates with each well containing coverslips. Cells were serum-starved (2% serum) 12 hrs before transfection. After 24 hrs, transfection was performed with either *BMP7* WT and MUT constructs (1µg). Forty-eight hrs of post-transfection, cells were washed with 1X PBS followed by immediate fixation with 4% paraformaldehyde (PFA) for 15 minutes. The fixed cells were incubated for 30 minutes with 0.5% Triton-X (Sigma-Aldrich, USA) prepared in 1X PBS for permeabilization, followed by blocking with 5% non-fat dry milk for 2 hrs at RT and then incubation with primary antibody.

##### Staining with BMP7 antibodies

The cells were incubated with 1:100 diluted BMP7 specific primary antibody (R&D systems Inc., USA) overnight at 4°C followed by 1X PBST washing (5 times) and incubation with goat anti-mouse Alexa Fluor 488 IgG (Molecular Probes, USA) for 2 hrs at RT. Then cells were washed with 1X PBST (5 times) and nuclei were stained with DAPI (Sigma-Aldrich, USA) and cytoskeleton with Phalloidin (Sigma-Aldrich, USA), mounted in mounting media and scanning was carried out using confocal microscope (Leica SP8 STED Laser Scanning Super Resolution Microscope System, Germany). The images were analysed by ImageJ software and assembled by Adobe Photoshop software.

##### Co-localization of BMP7 with ER-Tracker

In another independent experiments, colocalization of BMP7 with ER-Tracker (Invitrogen^TM^, USA) was performed in H9c2 cell lines following the afore-mentioned protocol. After the treatment of secondary antibody for BMP7, cells were washed with 1X PBST (5 times) and stained with ER-Tracker (Invitrogen^TM^, USA) to localise ER. The nuclei were strained with DAPI (Sigma-Aldrich, USA). After washing with 1X PBST (5 times) fixed cells were mounted in mounting media and confocal imaging was performed with Leica SP8 STED Laser Scanning Super Resolution confocal microscopy. The images were analysed by ImageJ and the statistical analysis was performed using LAS X software and images were assembled by Adobe Photoshop software.

##### Co-localization of BMP7 with SERCA2

Colocalization of BMP7 with SERCA2 was carried out in independent experiments following similar protocol. After treatment with BMP7 primary and secondary antibodies, the cells were washed with 1X PBST (3 times) blocked with 5% non-fat milk for 2 hrs at RT. Then the cells were incubated with diluted (1:100) SERCA2 primary antibody (Novus Biological, USA) overnight at 4°C. Following washing with 1X PBST washing (3 times), cells were incubated with goat-anti mouse IgG Alexa Fluor 568 (Molecular Probes, USA) at 1:100 for 2 hrs at RT. The cells were further washed with 1X PBST (5 times) and nuclei were stained with DAPI. Next, after washing with 1X PBST, the slides were mounted. The scanning was performed with confocal (Leica SP8 S TED Laser Scanning Super Resolution confocal microscopy), followed by image analysis by ImageJ and statistical analysis by LAS X software and assembly by Adobe Photoshop software.

##### Cell proliferation assay

EdU (5-ethynyl-2′-deoxyuridine) staining was performed to check cell proliferation. H9c2 cells were grown on glass cover-slip in 6-well culture plates. After 18 hrs, EdU (Invitrogen^TM^, USA) was added and transfection was performed as described above after 6 hrs of EdU addition. Cells were fixed with 4% PFA after washing with 1X PBS post 48 hrs of transfection. Fixed cells were washed twice with 3% BSA in PBS followed by permealization with 0.5% Triton-X. Following washing with 3% BSA, Click-iT reaction cocktail (Invitrogen^TM^, USA) was added and incubated for 1 hr at RT. The cells were washed twice with 3% BSA and nuclei were stained with DAPI followed by 1X PBS washing and mounting. The scanning was performed with confocal microscope (Zeiss LSM 780) followed by image analysis by ImageJ and assembly by Adobe Photoshop software.

#### 2.4.4 Estimation of the expression of BMP7 protein by Western Blotting

To estimate the expression of BMP7 muteins against WT, Western blotting was performed in P19 cells. Cells were seeded in 6 well-plates 24 hrs prior to transfection. Twelve hrs before transfection, cells were serum-starved. At 30-40% confluency, cells were transfected with *BMP7* WT and MUT constructs (1µg) followed by lysate preparation in RIPA Buffer (as per Dixit et al., 2021) after 48 hrs of transfection. Quantification of total proteins was performed by Bradford Assay. 25µg of non-reducing samples were diluted in Laemmeli’s loading buffer followed by denaturation for 5 min at 95°C and then lysates were resolved on 8% SDS polyacrylamide gel and proteins were transferred to PVDF membrane (Bio-Rad Laboratories Inc, CA, USA). The membrane was blocked using 5% dry skimmed milk in 1X TBST (25 mM Tris-Cl pH = 7.5, 0.15 M NaCl with 0.1% Tween 20) for 2 hrs at RT followed by incubation with BMP7-specific primary antibody (R&D systems Inc., USA), overnight at 4°C. Further, the membrane was washed five times (5 min each wash) in 1X TBST, followed by incubation with HRP conjugated goat anti-Mouse IgG antibody (Genei, Merck Specialties Pvt. Ltd. Germany) for 2 hrs at RT. The membrane was washed in 1X TBST ten times (5 min each wash) and visualized with enhanced chemiluminescence detection kit (Amersham, GE health Care, USA).

##### Estimation of SMAD1/5 Phosphorylation

To estimate the expression of phospho-SMAD1/5 (Ser463/465), cells were transfected with *BMP7* WT and MUT constructs (1µg) and lysate was prepared after 48 hrs of transfection. Similarly, the synergistic effect of *ALK2* was also checked by co-transfecting *ALK2* (1ug) along with *BMP7* WT and MUT constructs (1µg each) followed by lysate preparation and quantification. 25µg of each sample in Laemmeli’s loading buffer (reducing) were resolved on 8% SDS polyacrylamide gel followed by transferring the protein samples to PVDF membrane. Membrane was blocked in 5% non-fat dry milk in 1X TBST followed by overnight incubation with phospho-SMAD1/5 primary antibody (CST, USA) at 4°C subsequently washed and probed with HRP conjugated goat anti-Rabbit IgG secondary antibody (Genei, Germany) for 2 hrs at RT. Finally, the expression of phospho-SMAD1/5 was detected with chemiluminescence detection kit (GE health Care, USA) using chemiDoc (Amersham^TM^ Imager 680, Japan) following thorough washing (5 times, 5 min each in 1X TBST).

#### 2.4.5 Transactivation assay by Dual Luciferase Reporter Genes

*Dual luciferase reporter assay* was performed to study the impact of BMP7 muteins on the actication of downstream promoters *viz. Id1-luc*, *Id3-luc Tlx2-luc*, and *p(SBE)_4_-luc*. P19 cells were seeded in 24-well culture plates with cell density of 10^5^ cells per well. Transfection was performed (at 30-40% confluency) with *BMP7* WT and MUT constructs (125ng) along with each reporter plasmids (125ng) for independent experiments. A constitutively expressing Renilla luciferase plasmid, pRL-TK vector (25ng) (Promega, USA) was also co-transfected as an internal control plasmid. In co-transfection experiments with *ALK2* (125ng) (which is one of the active receptors of BMP-ligands) were also performed for each aforementioned promoters with *BMP7* constructs in independent sets of experiment. Forty-eight hrs post transfection, cells were washed with 1X PBS followed by lysates preparation in 1X passive lysis buffer (PLB). Dual-Luciferase Reporter Assay System, (Promega Corp, WI, USA) was used to measure luciferase activity (as per manufacturer’s instructions) using Synergy/ HTX multi-mode reader (BioTek Instruments, Inc., USA). Three independent experiments were performed in triplicate for each sample (WT and MUTs), using all four reporter plasmids and the data acquired were pooled and plotted as mean fold change with standard error of mean. Reporter plasmids used in luciferase reporter assays were obtained as a kind gift (Tlx2-luc from Prof. Jeffrey L. Wrana, University of Toronto, Ontario, Canada; *p(SBE)_4_-luc* from Dr. Peter ten Dijke, Netherlands; Id1-luc from Dr. Maria Genander, Karolinska Institutet, Stockholm, Sweden; and Id3-luc from Prof. Daniel J. Bernard, Quebec, Canada) to Prof. B. Mohapatra.

#### 2.4.6 Quantification using real-time PCR (qPCR) Assay

For evaluating the effect of BMP7 WT and MUT constructs on the expression level of downstream target genes such as *Smad1, Smad4, Smad5, Gata4, Nkx2.5, Irx4* and *Mef2c* along with *Bmp2, Bmp7* and left right patterning genes *Pitx2* and *Nodal* real time PCR assay was performed. P19 cells were seeded in 6-well culture plates and after 24 hrs, transfected with WT and MUT constructs (1µg) at 50% confluency. Forty-eight hrs post transfection, total cellular RNA was extracted using TRI Reagent (Merck, Germany) by following manufacturer’s protocol. The integrity of RNA was checked on 1% agarose gel and further quantified by using NanoDrop (ThermoFisher Scientific, USA). All extracted RNA were treated with DNase I (Thermo Fisher Scientific, USA) and 2 µg of DNase I-digested RNA was used to prepare cDNA using Revertaid First Strand cDNA synthesis kit (ThermoFisher Scientific, USA) and Random hexamers. The q-PCR assay was performed in triplicate in real-time PCR machine (QuantStudio 5, Applied Biosystems, CA, USA) using SYBR green (Sigma-Aldrich, USA) as per manufacturer’s instruction.

##### Statistical Analysis

One-way ANOVA test was performed using Graph pad Instat3 software (GraphPad Software, La Jolla California USA) for analysing western blotting, luciferase assay and real time PCR data followed by Dunnett’s post-hoc test. *p value* of <0.05 was considered statistically significant. Statistical analysis for colocalization was performed using Leica Microsystems software.

## 3. Results

### 3.1 Identification of genetic variants in *BMP7*

Sanger sequencing of the coding exons and the flanking exon-intron boundary regions of *BMP7,* in 285 CHD probands yielded 5 rare non-synonymous, 5 synonymous and six intronic variants. In addition, we detected 3 novel variations in 5’ UTR. The five missense variants namely c.254A>T; p.D85V, c.523C>T; p.R175W, c.847G>A; p.A283T, c.945G >A; p.M315I, c .962A>G; p.N321S are detected in 5 unrelated cases, in heterozygous condition with minor allele frequency (MAF) <0.01 **(Table 1)**. Screening of 400 ethnically matched healthy controls (800 chromosomes) did not reveal any of these variants. Four out of five variants are novel and submitted in ClinVar database. The allotted RCV numbers are., SCV002567819 (c.254A>T; p.D85V), SCV002567820 (c.523C>T; p.R175W), SCV002567821 (c.847G>A; p.A283T), SCV002567822 (c.945G >A; p.M315I). The p.N321S variant has been previously reported (rs767011450). The variants p.D85V and p.R175W are identified in TGF-β pro-peptide domain while p.A283T lies close to the cleavage site, and p.M315I and p.N321S are in mature peptide domain **Figure 1(A)**. The sequence chromatogram showing abovementioned heterozygous variations, its location on the protein domains and their conservation across the species are shown in **Figure 1(B)**.

**Figure 1.**
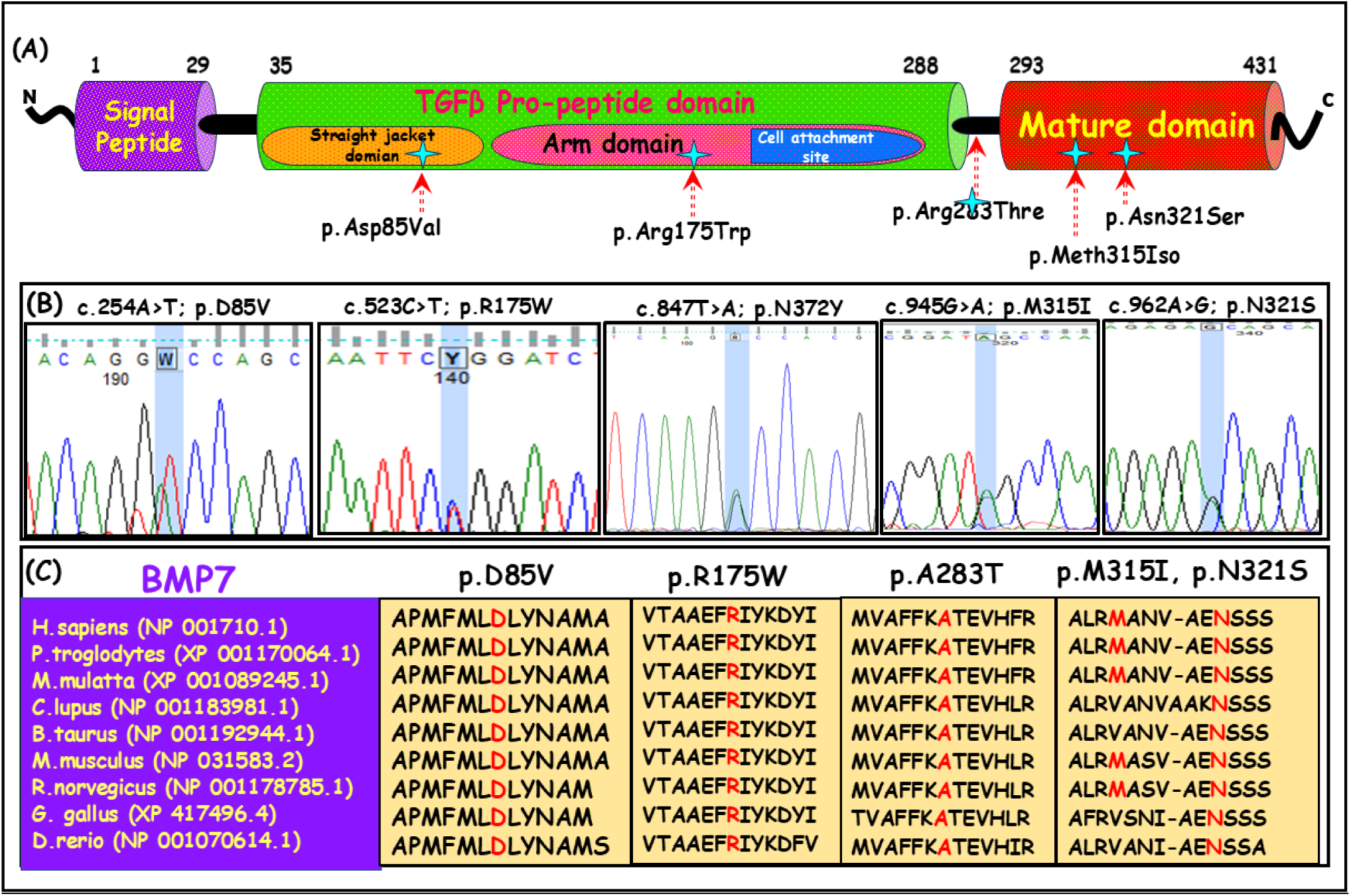
Schematic representation of BMP7 protein with identified variants and phylogenetic conservation analysis. **(A)** Diagrammatic representation of BMP7 protein with 431 amino acids showing different domains namely., signal peptide domain (1-29aa), TGFβ-propeptide domain (35-288aa), protease cleavage site (289-292) and mature peptide domain (294-431aa). Domain-wise location of different identified missense variants (marked with star). **(B)** The sequence chromatogram for the identified variants and cross species alignment of the substituted amino acid across different vertebrate species (highlighted in red) for all the five variants (p.D85V, p.R175W, p.A283T, p.M315I and p.N321S). **(C)** evolutionary conservation of mutated amino acid (highlighted in red) across different species of vertebrate for all the five variants (p.D85V, p.R175W, p.A283T, p.M315I and p.N321S).

**Table 1.** List of missense variants identified in BMP7 and its associated CHD phenotype with novelty status.

| Nucleotide change | Amino Acid change | CHD Phenotype | Patient ID | Parental genotype | | MAF | | BMP7 Domain | ClinVar | Indigenome | 1000 Genome database | Genome Asia100K | gnomAD | I-mutant 2.0 ($\Delta\Delta G$, RI) |
| --- | --- | --- | --- | --- | --- | --- | --- | --- | --- | --- | --- | --- | --- | --- |
|  |  |  |  | Father | Mother | Cases (N=285) | Controls (N=400) |  |  |  |  |  |  |  |
| c.254A>T | p.D85V | VSD | CHD356 | NA | NA | 1<br>(0.00175) | 0 | Pro-peptide | Novel<br>(SCV002567819) | NR | NR | NR | NR | Increase (0.79, 4) |
| c.523C>T | p.R175W | ASD, VSD | CHD1 | NA | CC | 1<br>(0.00175) | 0 | Pro-peptide | Novel<br>(SCV002567820) | NR | NR | NR | NR | Decrease (-0.82, 6) |
| c.847G>A | p.A283T | ToF | CHD881 | GA | GG | 1<br>(0.00175) | 0 | Pro-peptide | Novel<br>(SCV002567821) | NR | NR | NR | NR | Decrease (-1.03, 8) |
| c.945G>A | p.M315I | TAPVR, PA, ASD, IVC | CHD608 | NA | NA | 1<br>(0.00175) | 0 | Mature domain | Novel<br>(SCV002567822) | NR | NR | NR | NR | Decrease (-0.82, 6) |
| c.962A>G | p.N321S | PS, TAPVR | CHD657 | NA | NA | 1<br>(0.00175) | 0 | Mature domain | rs767011450 | 0.0229 | 0.008 | 0.0078 | Reported | Decrease (-1.40, 8) |

**Abbreviations:** VSD- Ventricular septal defects, ASD- Atrial septal defects, ToF- Tetralogy of Fallot, TAPVR- Total Anomalous Pulmonary Venous Return, PA- Pulmonary Atresia, IVC- Inferior vena cava, PS- Pulmonary stenosis, MAF- Minor allele frequency, dbSNP- single nucleotide polymorphism database, NR- Not reported, RI- Reliability index

### 3.2 Genotype-phenotype correlation

The novel variant p.D85V was detected in 6.5yrs old female diagnosed with VSD. Similarly, variant p.R175W that lies in TGFβ pro-peptide domain, was observed in 3 months old male infant affected with both ASD and VSD. The variant p.A283T which was localized in cleavage site, was detected in 1.5yrs male patient with ToF phenotype. A female child affected with complex CHD phenotypes, such as TAPVR, PA, ASD and IVC, was found to harbour p.M315I variant in mature peptide domain. Another mature peptide variant, p.N321S, was also identified in a male proband with complex CHD (PS, TAPVR). The parental genotype for the variant p.R175W showed that mother harbouring homozygous WT allele (CC). Paternal DNA sample was not available. For the variant p.A283T, the mother was homozygous wild-type (GG). However, the genotype of father was heterozygous MUT (GA). Due to unavailability of DNA samples, parental genotype of other variants could not be determined.

In addition to this, another novel variant c.1114>T; p.N372Y was also identified in 11 CHD probands exhibiting varied phenotype namely ASD (4), VSD (4), Dextrocardia (2), PS (1) and BAV (1). The variant was also found in 7 controls. Though the variant was novel, its frequency in the population was quite high. Therefore, further functional analyses were not performed for this variant.

### 3.3 Analysis of evolutionary conservation of BMP7 protein across species

Multiple sequence alignment of BMP7 protein across several vertebrate species showed amino acid residues at Asp85, Arg175, Ala283, and Asn321 are highly conserved. However, the amino acid residue Meth315 is conserved only in few mammalian species namely, *Homo sapiens, Pan troglodytes Macaca mulatta, Mus musculus*, and *Rattus norvegicus*. The conserved amino acids along with the flanking amino acid sequences has been presented in **Figure 1(C)**.

### 3.4 Potential pathogenicity of *BMP7* missense variants

The disease-causing potential of the identified variants was predicted by Varcard2 (http://www.genemed.tech/varcards2/#/index/home) that incorporate prediction of more than 15 bioinformatic tools. All the five missense variants p.D85V, p.R175W, p.A283T, p.M315I, p.N321S were predicted to be disease-causing by Mutation Taster, and damaging by FATHMM_MKL, M_CAP, CADD and Eigen. However, the prediction of Polyphen2_HVAR, PROVEAN, MetaSVM, MetaLR revealed that the pro-peptide domain variants (p.D85V, p.R175W) were probably damaging while rest of the mature domain variants (p.M315I, p.N321S) were benign. The prediction and the algorithmic scores of these variants are presented in **Table 2**.

**Table 2.**
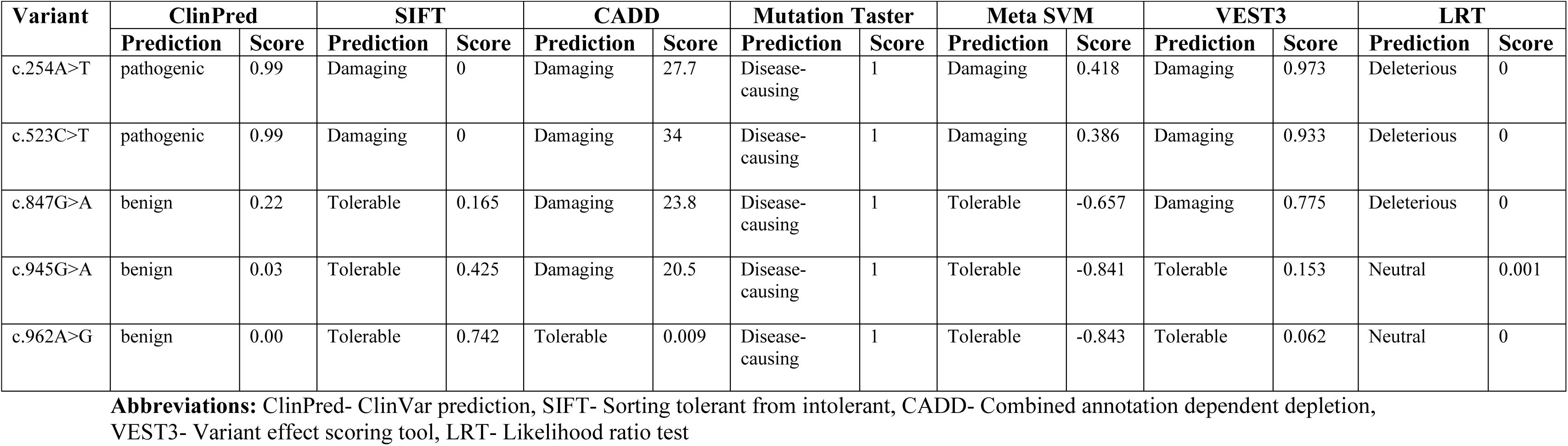
Table representing the in-silico analysis for predicting the disease-causing potential of BMP7 variants.

### 3.5 Changes in predicted secondary and tertiary structures of BMP7 muteins

Prediction of the changes in the secondary structures of BMP7 muteins, revealed that there were considerable changes in mutein, p.D85V, p.R175W, p.A283T, p.M315I and p.N321S **(Supplementary Figure 1 and Supplementary Table S2)**.

The 3-D modelling structures of BMP7 WT and muteins revealed comparable changes in the conformation of muteins as evident by superimposition of the WT and muteins structures by PyMOL. In prediction of protein structures, the global RMSD values at alpha carbon depicts the quantitative estimation of similarity or dissimilarity between two or more structures of proteins. The global RMSD value for the superimposed structures p.D85V versus BMP7 WT was calculated to be 3.016 Å which indicate relatively significant deformities in the tertiary structure of mutein. Similarly, considerable changes in the 3-D structures of other muteins (p.R175W, p.A283T, p.M315I and p.N321S) were visualized with supported RMSD value of 1.682 Å, 3.501 Å, 3.568 Å and 5.780 Å, respectively which suggested that even a single amino acid change could alter the conformation of proteins greatly. The comparable superimposed structures of WT versus each muteins were presented in **(Figure 2).**

**Figure 2.**
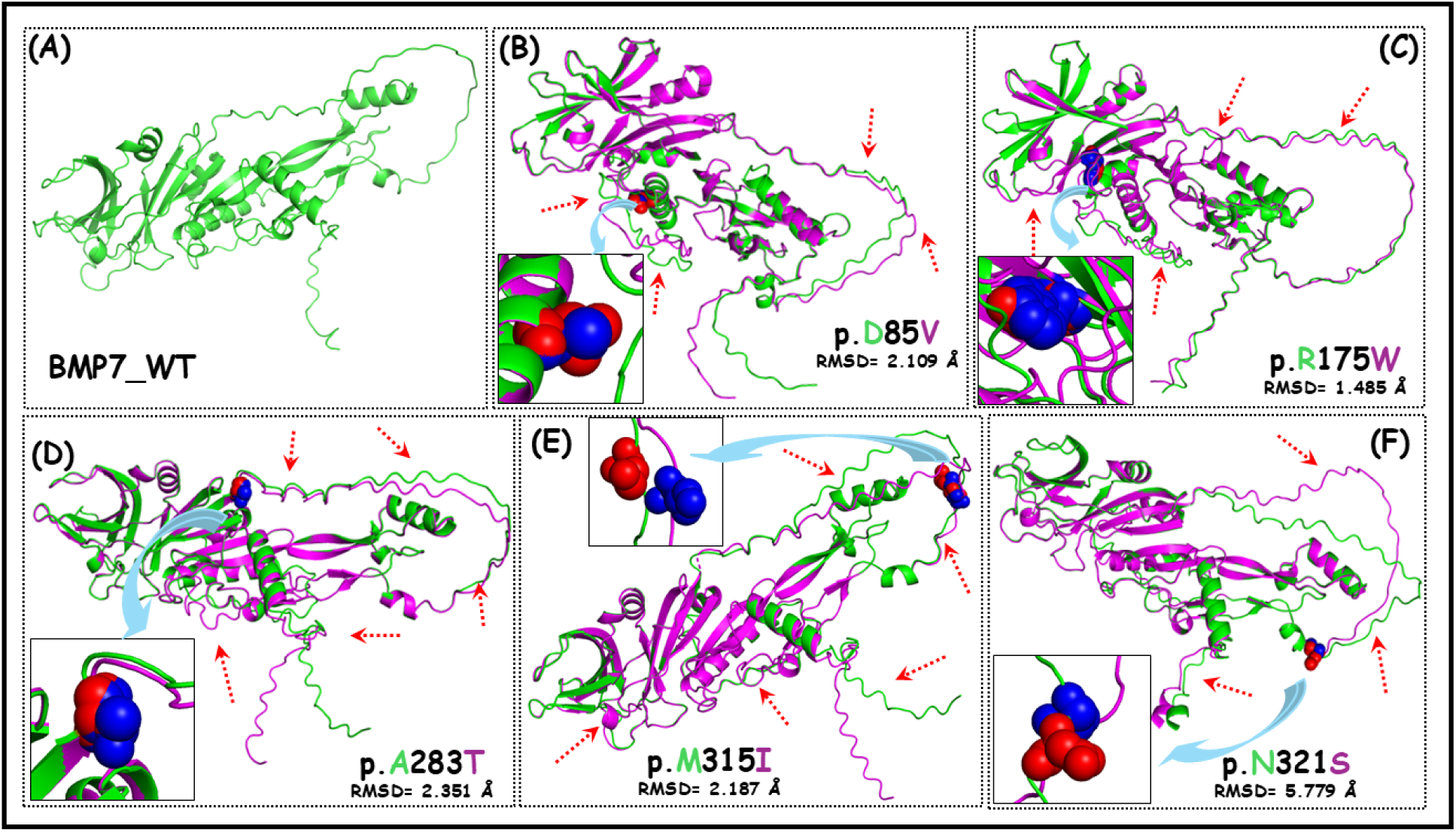
**(A-F)** Tertiary structures comparison of BMP7 WT and muteins. **(A)** modelled-tertiary structure of BMP7 WT, **(B-E)** superimposed tertiary structures of different muteins of BMP2 (p.Asp85Val, p.Arg175Trp, p.Ala283Thr, Met315Iso and p.Asn321Ser) over WT with supported global alpha carbon RMSD values. The tertiary structure of WT is shown in green colour while muteins are represented in magenta with mutated residues in spherical form. The conformational changes in the structures of muteins are indicated by red arrows.

### 3.6 Effect of variants on the expression of BMP7 protein and cellular localization

To check the cellular localization of the BMP7-WT and MUT proteins, immunofluorescence staining was also performed in H9c2 cells. The result demonstrated that the WT as well as all Muteins were localized in cytoplasm (**Figure 3(A1-F4))**. Since, BMP7 is a secretory protein which is likely transported through endoplasmic reticulum via Golgi to extracellular matrix, (Sedlmeier and Sleeman, 2017) we presumed that these proteins might be localized to endoplasmic reticulum (ER). To check trafficking process and effect of mutant on it, colocalization experiment with SERCA2 was performed which is an ER-specific protein. BMP7 WT and all MUTs were found to co-localize with SERCA2 **(Figure 3(A5-F8))**. Similarly, in another experiment, BMP7 WT and all muteins were also observed to co-localize with ER-Tracker **(Figure 3(A9-A12))** confirming their accumulation in ER in both the experiments. The quantitative analysis of colocalization was performed using Leica Application Suite X (LAS X) software, along with the colocalization scattered plot. Mander’s overlap coefficient (MOC) for colocalization with SERCA2 showed strong positive corelation scores viz. 0.8383, 0.8759, 0.8138, 0.8295, 0.73 and 0.8266 and also a positive Pearson’s correlation coefficient (PCC) scores of 0.7144, 0.7988, 0.7278, 0.6747, 0.5638 and 0.8015 due to BMP7 WT, p.D85V, p.R175W, p.A283T, p.M315I and p.N321S respectively. Similarly, the MOC scores for colocalization with ER-Tracker was 0.5933, 0.6881, 0.6508, 0.7671, 0.6667 and 0.6522 and the Pearson’s CC was measured to be 0.5045, 0.552, 0.5454, 0.7031, 0.5726 and 0.6087 in case of BMP7 WT, p.D85V, p.R175W, p.A283T, p.M315I and p.N321S respectively **(Figure 3(A9-F12))** which indicate that all the BMP7 variants affect the trafficking process significantly with more positively colocalized with ER.

**Figure 3.**
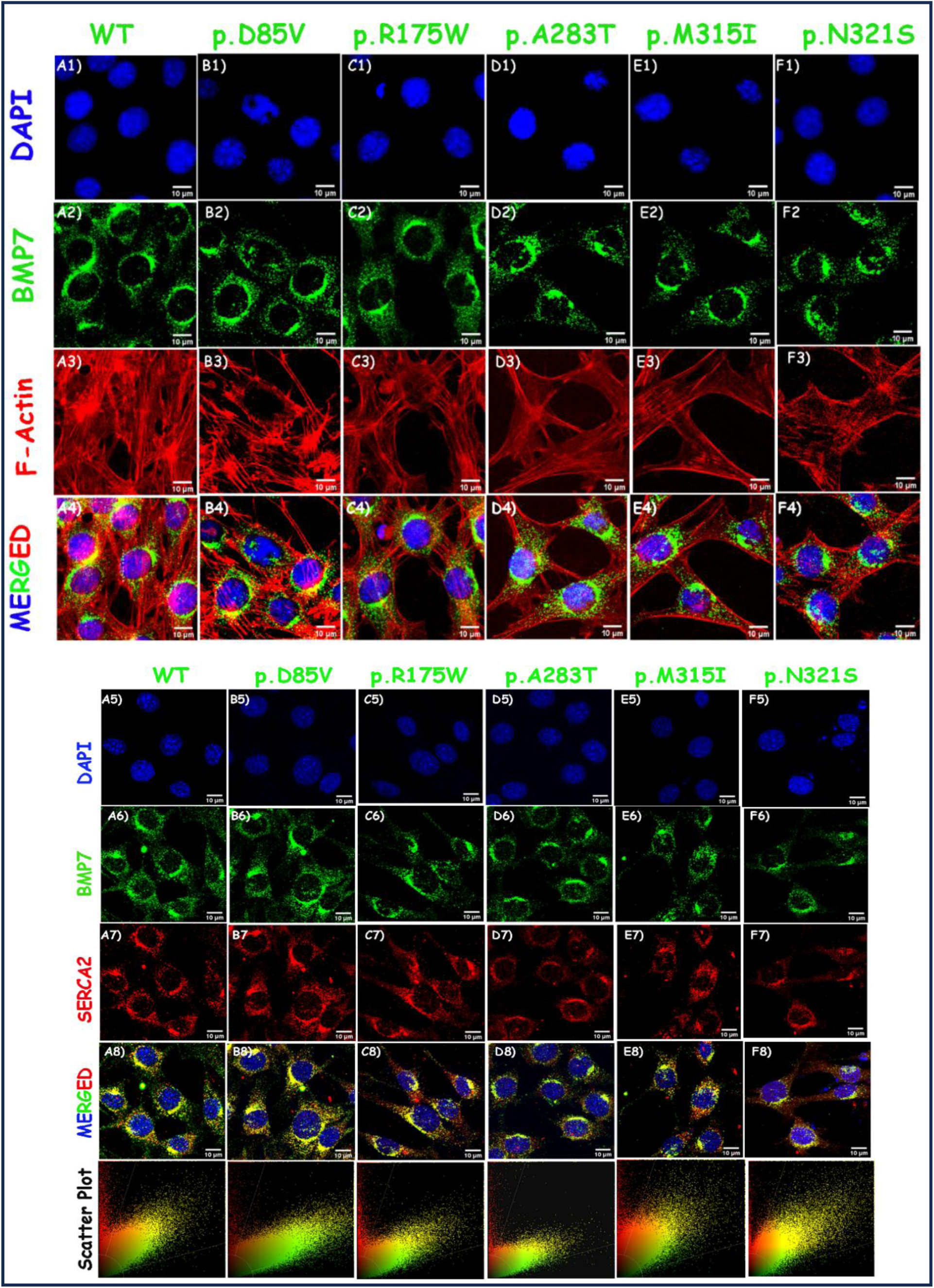
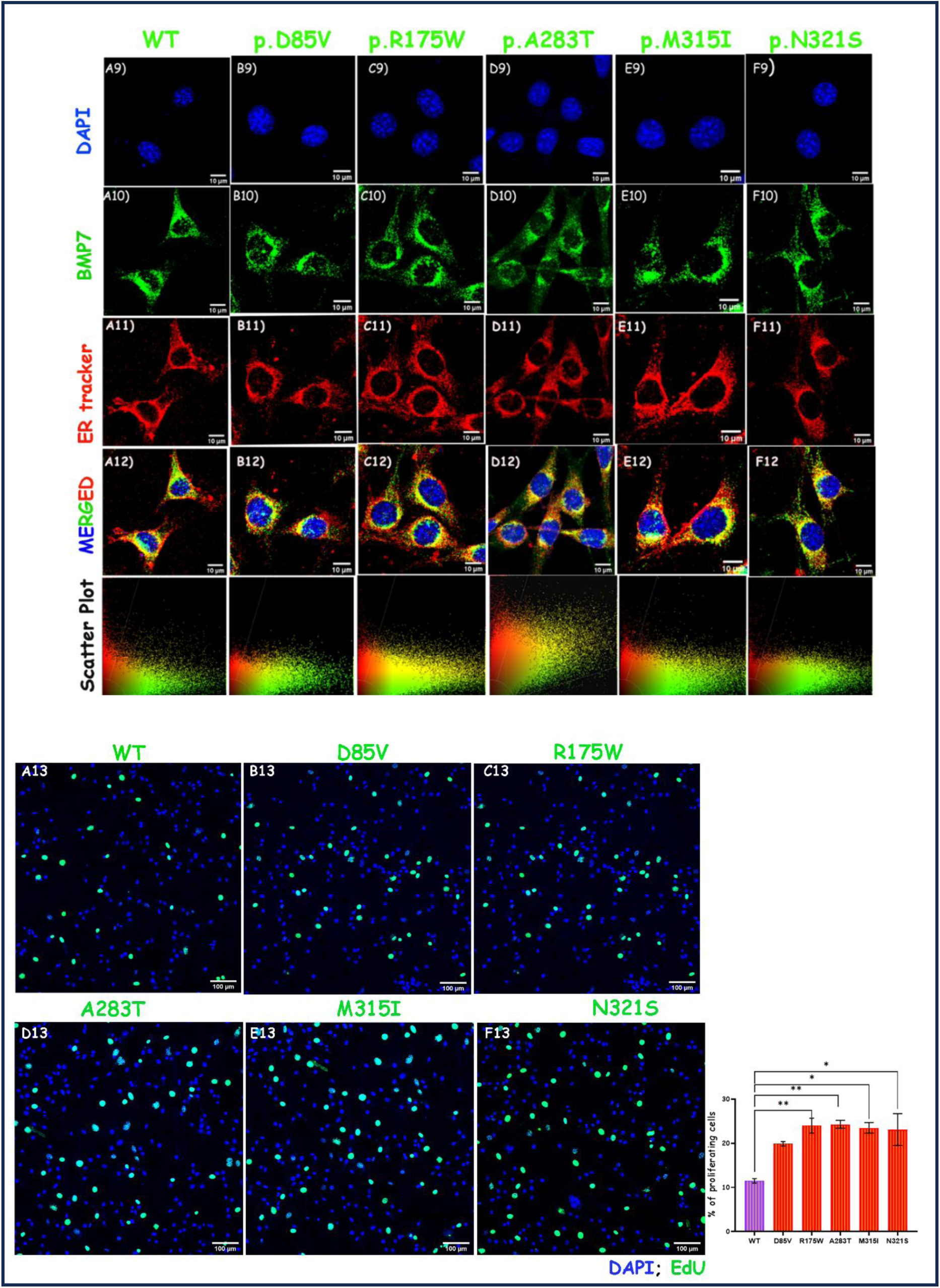
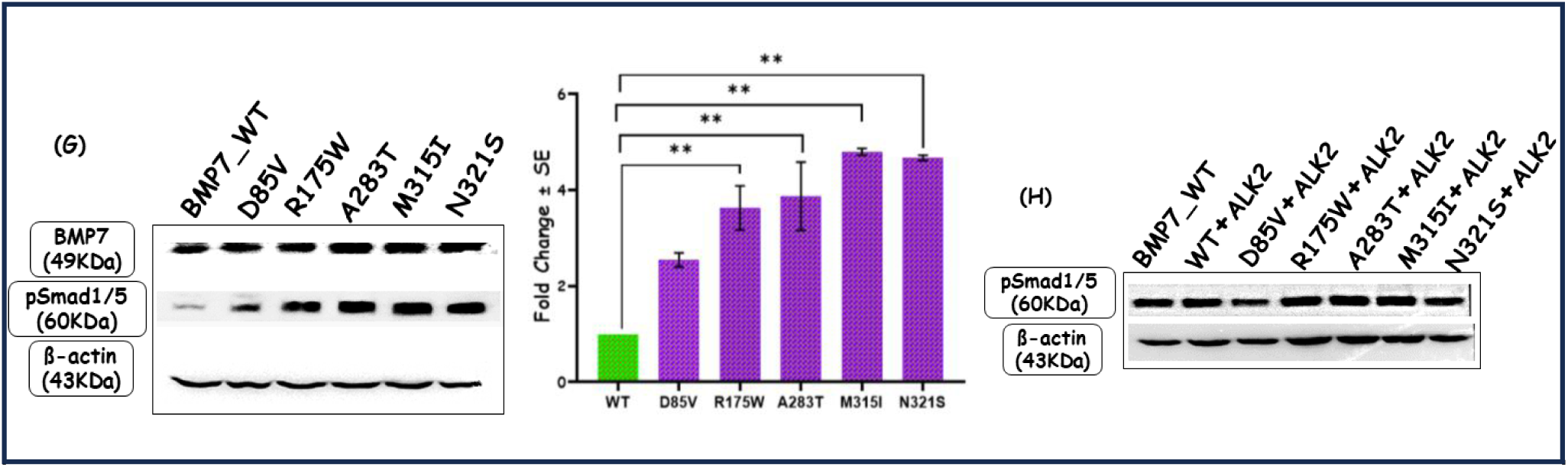
Expression analysis of BMP7 wild-type (WT) and muteins (MUT) by immunostaining and immunoblotting and colocalization of BMP7 proteins with SERCA2 and ER-Tracker. **(A1-F4)** Immunostaining in H9c2 cells showing cellular location of BMP7 WT and MUT (p.D85V, p.R175W, p.A283T, p.M315I and p.N321S). **(A5-F8)** Immunostaining in H9c2 cells showing colocalization of BMP7 proteins with SERCA2 with quantitative assessment by Scatter plot. **(A9-A12)** Immunostaining in H9c2 cells showing colocalization of BMP7 proteins with ER-Tracker with quantitative assessment by Scatter plot. **(A13-F13)** EdU staining showing cell proliferation due to all the variants in H9c2 cells. **(G)** Western blot analysis of WT and MUT in P19 cells showing expression of BMP7 and pSMAD1/5 due to WT and all five variants (p.D85V, p.R175W, p.A283T, p.M315I and p.N321S) with graphical representation in terms of fold change for pSmad1/5. **(H)** Western blotting showing expression level of pSMAD1/5 due to WT and all five variants (p.D85V, p.R175W, p.A283T, p.M315I and p.N321S) when co-transfected with ALK2.

However, there was no significant change in expression of muteins (p.D85V, p.R175W, p.A283T, p.M315I and p.N321S observed when compared with the WT in immunostaining.

### 3.7 Impact of BMP7 variants on protein expression by Western blot

Western blot analysis of BMP7 proteins was performed to evaluate the expression level of BMP7 WT and muteins in P19 cells. No significant differences could be observed in the level of expression of BMP7, in cells expressing either WT or mutant proteins **(Figure 3(G))**. However, evaluation of the expression level of phosphor-SMAD1/5 showed significant differences. The expression level of phospho-SMAD1/5 was upregulated by 2.544 (p<0.0712), 3.627 (p<0.0068), 3.869 (p<0.0044), 4.795 (p<0.001), 4.669 (p<0.0012) fold due to all the five variants p.D85V, p.R175W, p.A283T, p.M315I and p.N321S respectively **(Figure 3(G))**. In contrast, the co-transfection of BMP7 with ALK2 did not cause any change in the expression level of pSMAD1/5 in case of BMP7 WT or all the five variants (p.D85V, p.R175W, p.A283T, p.M315I and p.N321S) **(Figure 3(H))**.

### 3.8 Effect of BMP7 variants on transactivation of downstream promoters

To evaluate the functional differences of muteins over the WT, dual reporter assay was performed *in vitro,* with cardiac specific downstream promoters of BMP signaling namel*y Id1-luc*, *Id3-luc*, *Tlx2-luc* and *p(SBE)_4_-luc.* The level of transactivation of all the promoters in response to either BMP7 WT or MUTs was normalized with the basal activity of *p(Id1)-*, *p(Id3)-*, *p(Tlx2)-* and *p(SBE)_4_-luc* promoters respectively. The fold change in the activity of these promoters in response to BMP7 MUTs was calculated by comparing with the WT. The synergistic activity of BMP7 was also estimated by co-transfection experiment with *ALK2* with each aforementioned promoter. The transactivation of all the promoters i.e., *Id1-luc* (3.85 fold, p<0.0031), *Id3-luc* (5.15 fold, p<0.0049), *Tlx2-luc* (3.07 fold, p<0.0001) and *p(SBE)_4_-luc* (4.75 fold, p<0.0001) **(Figures 4A, 4C, 4E & 4G]** by wild-type BMP7. Moreover, when BMP7 wild-type was co-transfected with *ALK2,* it synergistically further trans-activated *Id1* (3.4 fold, p<0.0001), *Id3* (1.79 fold, p<0.0001), *Tlx2* (1.78 fold, p<0.0001) and p(SBE)_4_-luc (1.44 fold, p<0.0002) compared to BMP7 alone **[Figures 4B, 4D, 4F &4H]**.

**Figure 4.**
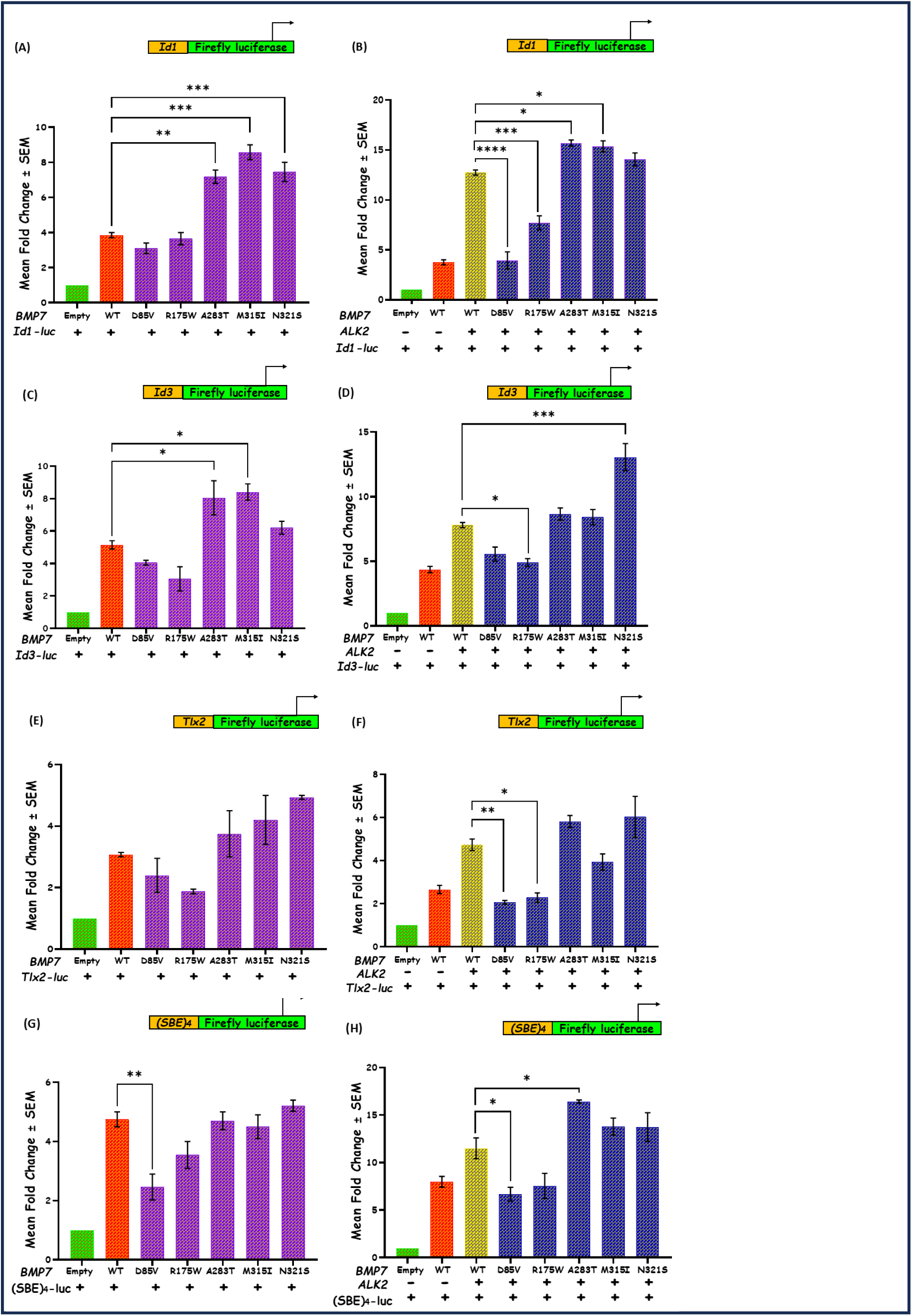
Dual luciferase reporter assay for measuring the transactivation of Id1-luc, Id3-luc, Tlx2-luc and p(SBE)_4_-luc promoters. (A) Transcriptional activity of Id1-luc due to BMP7 WT and MUT alone, (B) synergistic activity of Id1-luc due to WT and MUT when co-transfected with ALK2. (C) Transcriptional activity of Id3-luc due to BMP7 WT and MUT alone, (D) synergistic activity of Id3-luc due to WT and MUT when co-transfected with ALK2. (E) Transcriptional activity of Tlx2-luc due to BMP7 WT and MUT alone, (F) synergistic activity of Tlx2-luc due to WT and MUT when co-transfected with ALK2. (G) Transcriptional activity of p(SBE)_4_-luc due to BMP7 WT and MUT alone, (B) synergistic activity of p(SBE)_4_-luc due to WT and MUT when co-transfected with ALK2.

The pro-peptide domain variant p.D85V, caused marginal reduction in the expression of promoters *Id1-luc* (1.24 fold, p=0.515) **[Figure 4A]**, Id3 (1.27fold, p=ns) **[Figure 4C]** and *Tlx2-luc* (1.27 fold, p=0.8057) **[Figure 4E]**, while a significant reduction in case of p(SBE)_4_-luc promoter (1.93 fold, p=0.0071) **[Figure 4G]**. However, co-transfection of this variant with *ALK2* showed significant decrease in the activity of these promoters, *Id1-luc* (3.23 fold, p <0.0001), *Id3-luc* (1.4 fold, p=0.0708), *Tlx2-luc* (2.29 fold, p=0.0074) and *p(SBE)_4_-luc* (1.72 fold, p=0.0287) **[Figures 4B, 4D, 4F and 4H]**. Similarly, other pro-peptide domain variant, p.R175W also reflect the comparable trend of decrease in the activity of promoters, *Id1-luc* (1.05 fold, p=0.9956), *Id3-luc* (1.68 fold, p=0.1193), *Tlx2-luc* (1.6 fold, p=0.3633) and *p(SBE)_4_-luc* (1.34 fold, p=0.1308) **[Figure4A-G]**. Further, synergistic reduction in the activity of these promoters was also observed when co-transfected with *ALK2* by 1.65 fold (p=0.0007), 1.59fold (p=0.0205), 2.07 fold (p=0.0122), and 1.52 fold (p=0.0726) **[Figures 4B-H]** with *Id1-luc*, *Id3-luc*, *Tlx2-luc* and *p(SBE)_4_-luc* respectively.

In contrast to pro-peptide domain variants, the cleavage site variant, p.A283T show significant increase in expression of the promoters *Id1-luc* (1.86 fold, p=0.0012) **[Figure4A]**, *Id3-luc* (1.56 fold, p=0.0315) [Figure4c], *Tlx2-luc* (1.22 fold, p=0.7972) **[Figure4E]** and *p(SBE)_4_-luc* (1..01fold, p=0.9999) **[Figure 4G]**. Parallelly, the increased activity of these promoters was also observed by 1.23 fold (p=0.0188) **[Figure 4B]**, 1.11 fold (p=0.7607) **[Figure 4d]**, 1.23 fold (p=0.3435) **[Figure 4(F)]**, 1.42 fold (p=0.0264) **[Figure 4(H)]** respectively, when this variant was co-transfected with *ALK2*.

Likewise, the mature domain variant p.M315I showed significant increase in the transactivation of promoters *Id1-luc* (2.23 fold, p<0.0001) **[Figure 4A]**, *Id3-luc* (1.63 fold, p= 0.0181) **[Figure 4C]**, while two other promoters i.e., *Tlx2-luc* (1.36 fold, p=0.4088) **[Figure 4E]** and *p(SBE)_4_-luc* (1.05 fold, p=0.9824) **[Figure 4G]** though not significant, however showed an increasing trend. Further, with co-transfection of *ALK2,* synergistic transactivation of all the four promoters with a modest increasing trend was observed i.e., *Id1-luc* (1.21 fold, p=0.0346) **[Figure4B]**, *Id3-luc* (1.07 fold, p=0.9238) **[Figure 4D]**, *Tlx2-luc* (1.2 fold, p=0.6018) **[Figure 4F]**, and *p(SBE)_4_-luc* (1.19 fold, p=0.4155) **[Figure 4H]**. The effect of p.N321S, another variant in mature peptide domain, also demonstrated similar trend of increasing activity of all the promoters *Id1-luc* (1.94 fold, p< 0.0007) **[Figure 4A]**, *Id3-luc* (1.2 fold, p=0.6161) **[Figure 4C]**, *Tlx2-luc* (1.61 fold, p=0.0968) **[Figure 4E]** and *p(SBE)_4_-luc* (1.09 fold, p=0.8181) **[Figure 4G]**. Interestingly, the synergistic activity of all the promoters also demonstrated remarkable increase when co-transfected with *ALK2* by 1.1 fold (p=0.3997) **[Figure 4B]**, 1.67 fold (p< 0.0005) **[Figure 4D]**, 1.27 fold (p=0.2051) **[Figure 4F]**, and 1.19 fold (p=0.4356) **[Figure 4H]** with *Id1-luc*, *Id3-luc*, *Tlx2-luc* and *p(SBE)_4_-luc* respectively.

### 3.9 Effect of BMP7 variants on the expression of downstream target genes

Since the Western blot analysis of the BMP7 variants revealed no difference in the expression pattern of the gene, we further checked its expression pattern by qRT-PCR. The assay demonstrated, significant increase in the expression pattern of *Bmp7* variants: p.D85V (1.50 fold, p=0.002), p.R175W (1.51 fold, p=0.0014), and p.M315I (1.88 fold, p<0.0001). However, p.A283T (1.22 fold, p=0. 4070) and p.N321S (1.301, p=0.1295) showed only marginal enhancement in the expression of *Bmp7,* compared to WT **(Figure 5A)**. Interestingly, *Bmp2,* that forms a potent heterodimer with *Bmp7* and enhances its activity, also exhibited significant increase in its expression, in response to variant p.D85V (1.322 fold, p<0.0001), p.R175W (1.172 fold, p<0.0001), p.A283T (1.325 fold, p<0.0001), p.M315I (1.137 fold, p=0.0024) and p.N321S (1.261 fold, p<0.0001) **(Figure 5B)**.

**Figure 5.**
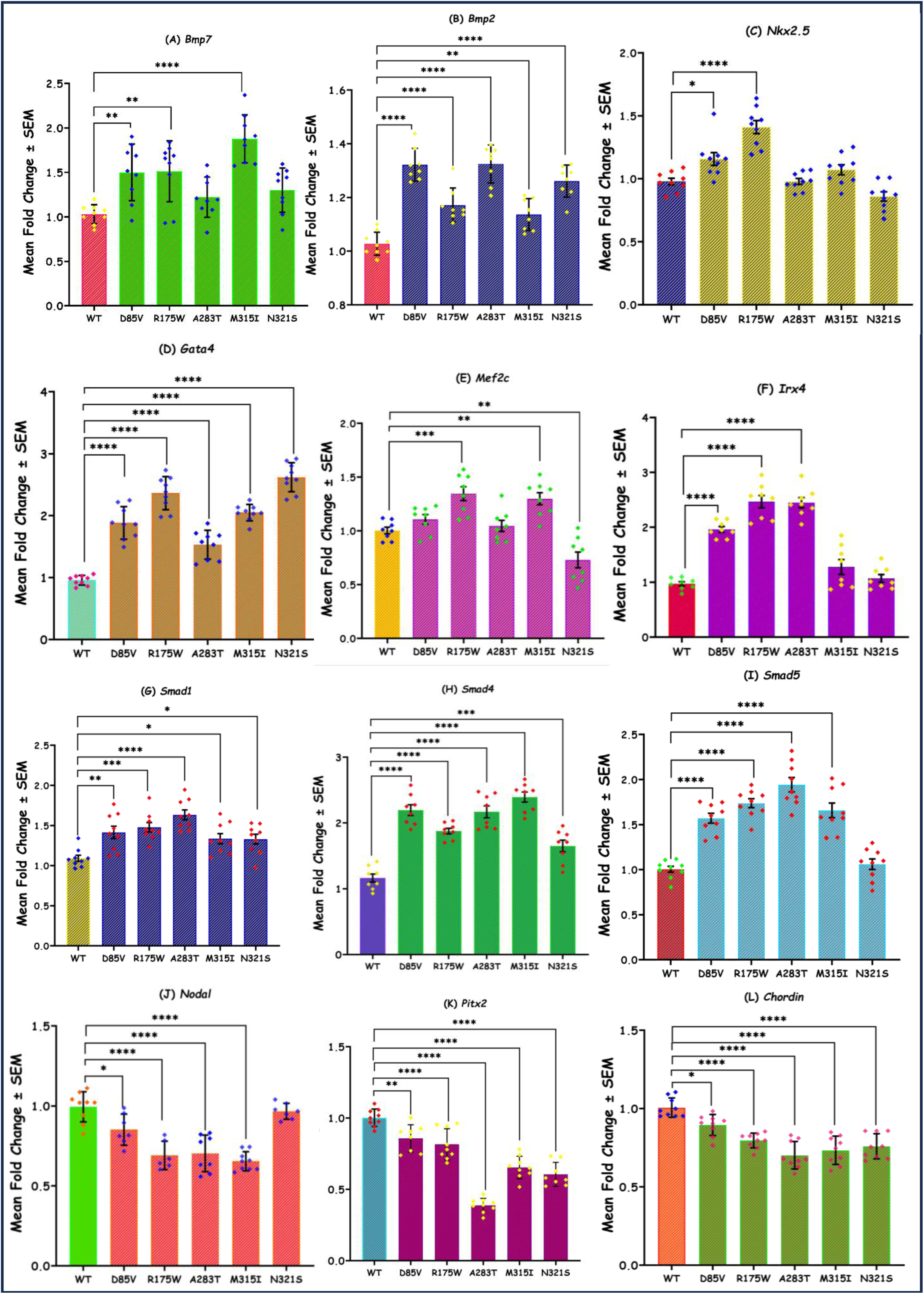
qRT-PCR analysis showing the expression of different downstream targets genes of BMP7 **(A)** Bmp7, **(B)** Bmp2, **(C)** Nkx2.5, **(D)** Gata4, **(E)** Mef2c, **(F)** Irx4, **(G)** Smad1, **(H)** Smad4, **(I)** Smad5, **(J)** Nodal, **(K)** Pitx2, **(L)** Chordin due to all the five variants (p.D85V, p.R175W, p.A283T, p.M315I and p.N321S) of BMP7.

We further checked the impact of BMP7 variants on the endogenous expression of certain other downstream target genes of BMPs. The cardiac enriched transcription factors such as ***Nkx2-5***, ***Gata4***, ***Mef2c*** and ***Irx4*** showed over all enhanced expression pattern. Up-regulation of ***Nkx2.5*** was also considerably significant due to pro-peptide domain variants p.D85V (1.16 fold, pc=0.0110), p.R175W (1.41 fold, p<0.0001) whereas the other variants p.A283T (0.9788 fold, p= >0.9999) p.M315I (1.07 fold, p=0.3326) and p.N321S (0.86 fold, p=0.1490) did not cause any significant impact on ***Nkx2.5*** expression **(Figure 5C)**. Similar increasing trend was also observed with transcription factor ***Gata4*** that exhibited 1.88 fold (p<0.0001), 2.36 fold (p<0.0001), 1.53 fold (p<0.0001), 2.05 fold (p<0.0001), 2.62 fold (p<0.0001), in response to variants p.D85V, p.R175W, p.A283T, p.M315I and p.N321S respectively **(Figure 5D)**. However, p.R175W (1.35 fold, p=0.0004) and p.M315I (1.3 fold, p=0.0024) variants caused significant enhancement in the expression of ***Mef2c*** transcription factor while p.N321S (0.7297 fold, p=0.0051) causes significant down-regulation and the remaining two variants p.D85V (1.11 fold, p=0.5561) and p.A283T (1.05 fold, p=0.9760) had no significant effect on the expression of ***Mef2c* (Figure 5E)**. Additionally, remarkable upregulation of ***Irx4* (Figure 5F)** due to these variations such as p.D85V (1.96 fold, p<0.0001), p.R175W (2.47 fold, p<0.0001), p.A283T (2.45 fold, p<0.0001), p.M315I (1.28 fold, p=0.0807) and p.N321S (1.07 fold, p=0.9018) was also observed.

When we analysed 3 other co-transcriptional activators, the expression of *Smad1*, *Smad4*, *Smad5* was grossly increased. In the case of ***Smad****1,* the two pro-domain and cleavage site variants i.e., p.D85V (1.41 fold, p=0.0006), p.R175W (1.48 fold, p=0.0002), and p.A283T (1.63 fold, p<0.0001) caused significant enhanced expression of *Smad1* while moderately significant increase was observed for the remaining two mature domain variants p.M315I (1.33 fold, p=0.0289) and p.N321S (1.33 fold, p=0.0340) **(Figure 5G)**. Further, the expression of ***Smad4* (Figure 5H)** was also significantly upregulated with fold change of 2.20 fold (p<0.0001), 1.88 fold (p<0.0001), 2.17 fold (p<0.0001), 2.39 fold (p<0.0001), 1.65 fold (p=0.0002) in response to all five variants respectively. Increase in expression of ***Smad5*** was also observed due to p.D85V (1.57 fold, p<0.0001), p.R175W (1.74 fold, p<0.0001), p.A283T(1.94 fold, p<0.0001), p.M315I(1.66 fold, p<0.0001) and p.N321S (1.06 fold, p=0.9506) variants respectively **(Figure 5I)**.

Likewise, 2 left-right patterning gene (*Nodal* and *Pitx2*) was also analysed by qRT-PCR assay. Interestingly, the expression of L/R patterning genes, ***Nodal*** was significantly reduced in response to p.D85V (0.8519 fold, p=0.0106), p.R175W (0.6905 fold, p<0.0001), p.A283T (0.7033 fold, p<0.0001), p.M315I (0.6544 fold, p<0.0001), and p.N321S (0.9668 fold, p=0.9599) variants **(Figure 5J)**. Similar trend was observed in the expression of ***Pitx2*** where the expression was also significantly decreased due to all the five variants p.D85V (0.8573 fold, p=0.0031), p.R175W (0.8160 fold, p<0.0001), p.A283T (0.3882 fold, p<0.0001), p.M315I (0.6537 fold, p<0.0001) and p.N321S (0.6051 fold, p<0.0001) (**[Figure 5K)**. Moreover, significant downregulation of ***Chordin*** (an antagonist of BMPs) was caused due to p.D85V (0.8951 fold, p=0.0115) p.R175W (0.7962 fold, p<0.0001), p.A283T (0.7019 fold, p<0.0001), p.M315I (0.7334 fold, p<0.0001) and p.N321S (0.7593 fold, p<0.0001) variants (Figure 5L).

### 3.10 Effect of BMP7 mutants on cell proliferataion

Our data from protein expression (WB) study, as well as luciferase reporter assay and RT-PCR data have showed that the BMP7 mutants have GoF. In order to validate this, cell proliferation assay was performed as one of the study by Bongiovanni et al., 2024 demonstrated that BMP7 GoF results in increased cardiomyocytes proliferation (Bongiovanni *et al*., 2024). Hence, to check cell proliferation, EdU staining was performed with BMP7 WT and MUTS. The result showed that the mitotic index increased by 8.4% (p=0.0509), 12.5% (p=0.0086), 12.8% (p=0.0077), 12% (p=0.0105) and 11.6% (p=0.0124) due to p.D85V, p.R175W, p.A283T, p.M315I and p.N321S variants respectively when compared with BMP7 WT **(Figure 3A13-F13)**.

## 4. Discussion

BMP7 (chr. location: 20q13.31, HGNC: 1074), is a secretory protein, belonging to TGF-β superfamily, which gets translated into a 431 amino acid long protein (46.12KDa) that possesses a signal peptide (29aa), a pro-peptide domain (254aa), and a mature peptide domain (139aa). The signal peptide assists in directing the mature peptide to secretory pathway via endoplasmic reticulum (von Heijne, 1990), the pro-domain (PD) after hydrolysis by proteases, assists in regulating the stability and the dimerization of mature peptides (Sedlmeier and Sleeman, 2017; Moustakas and Heldin, 2009). The mature peptide forms functional homo/heterodimers via di-sulphide bonds, and secreted into extracellular matrix for initiation of BMP signalling (Hogan, 1996). During cardiogenesis, BMP7 is believed to play essential role during left-right patterning, epithelial to mesenchymal transformation (EMT) that led to endocardial cushion development followed by valvulogenesis (Kim *et al*., 2001) as well as in cardiomyocyte differentiation (Monzen *et al*., 2002). Furthermore, it also improves cardiac function in adult hearts following myocardial infarction by preventing pathological remodeling and fibrosis (Jin *et al*., 2018).

In the current investigation, we have identified five rare missense variants by screening 285-isolated CHD cases, none of which occurred in 400 control individuals (800 chromosomes). As per the genome databases on Indian (IndiGenome and and INDEX-db), Asian (Genome Asia, 1000 Genome and GnomAD) Populations, four out of these five variants are novel, the C-terminal variant, p.N321S (rs61733438), is reported in majority of databases, (MAF <0.01) though not found in any of the controls, examined by us. However, this variant has been previously identified in 3 patients with congenital eye defects (Wyatt *et al*., 2010). Two missense variants (p.D85V and p.R175W) which are confined to pro-peptide domain are associated with minor CHD phenotype i.e., VSD and ASD-VSD respectively. The variant p.A283T that lie close to the cleavage site, where furin-like protease split the protein, is identified in a proband with ToF. In addition, two other variants (p.M315I and p.N321S) in the mature peptide-domain, are detected in patients with severe CHD phenotype i.e, TAPVR, PA, ASD, IVC and PS, TAPVR respectively. To the best of our knowledge, this is the first case-control study of *BMP7* gene, where clinically diverse forms of CHDs are associated with variants in *BMP7*. Previously, *BMP7* mutations have been associated with several other conditions viz., in cases of hypospadias (Chen *et al*., 2007; Bouty *et al*., 2019), and variable ocular, kidney, brain, ear, palate, and skeletal anomalies. (Wyatt *et al*., 2010). *BMP7* pathogenic mutations have also been associated with combined pituitary hormone deficiency (Breitfeld *et al*., 2013). Furthermore, a list of polymorphisms in *BMP7* have been reported in bone mineralization and vascular calcification in the coronary, carotid, and abdominal aorta in a diabetes-enriched cohort of European (Freedman *et al*., 2009). Besides *BMP7*, pathogenic mutations causing CHD have been previously reported in other members of BMP family namely *BMP2* (Li *et al*., 2016; Ahluwalia and Gelb, 2021; Yogi *et al*., 2023), *BMP4* (Wang *et al*., 2014; Li *et al*., 2016; Posch *et al*., 2007; Qian *et al*., 2014) and *BMP10* (Dong *et al*., 2024). We have also identified several disease-causing mutations in *BMP2 a*nd *BMP4* in the same cohort (unpublished data). Besides these, CHD associated mutations in other members of BMP signaling have also been investigated previously. Loss of function mutations have been identified in *ALK2* (Joziasse *et al*., 2011), *ALK3* (Demal *et al*., 2019), *BMPRII* (Roberts *et al*., 2004; Qi *et al*., 2020), *SMAD1* (Wang *et al*., 2022), *SMAD4* (Wang *et al*., 2023), *GATA4* (Butler *et al*., 2010; Dixit *et al*., 2018), *NKX2.5* (McElhinney *et al*., 2003; Dixit *et al*., 2021), *MEF2C* (Qiao *et al*., 2017; Lu *et al*., 2018), *IRX4* (Cheng *et al*., 2011) in CHD. However, reports on association of CHD phenotype with BMP7 mutation are meagre.

BMP7, plays a critical role during embryogenesis. In a mouse model study, it has been shown that expression of BMP7 starts as early as gastrulation i.e., anterior primitive streak, node and axial mesoderm etc. and later on it has been detected in developing kidney, brain, eye, heart, tooth bud and intestine as well (Lyons *et al*., 1995; Helder *et al*., 1995). Further, it has been noted that BMP7 is expressed ubiquitously throughout the myocardium of the heart tube, including cardiac cushion (Lyons *et al*., 1995; Solloway and Robertson, 1999). In another study, Bmp7^R-GFlag^ homozygotes are seen to have a common atrium, a smaller right ventricle and a small, malformed OFT (Kim *et al*., 2019). The heterogenous phenotypes observed in this study, including valve, septa and OFT defects such as VSD; ASD, VSD; ToF; TAPVR, PA, ASD, IVC; PS, TAPVR are strongly associated with *BMP7* mutations.

The human BMP7 is a 431 aa long glycoprotein and the 29 aa long signal sequence target the protein to secretory pathway via endoplasmic reticulum (von Heijne, 1990). In this study, immunostaining of BMP7 clearly demonstrated cytoplasmic localization. Further, co-immunostaining of BMP7 with either SERCA2 (Sarcoplasmic Reticulum Ca^2+^-ATPase) or ER-Tracker distinctly demonstrated colocalization of BMP7 with both of these ER-markers, thus confirming BMP7 trafficking via ER-Golgi transport pathway. Moreover, both WT as well as mutant proteins (muteins) showed similar colocalization pattern. Although there was no significant change in expression pattern, muteins exhibit more positive colocalization with ER-tracker as analysed by both MOC and PCC, which is presumed to be due to possible delay in transport. Similar trafficking defect has also been reported in a mature domain mutant, p.V407I, in BMP10 (Hirono *et al*., 2019).

Out of 5 rare variations detected in this study, two (p.D85V and p.R175W) are located in the pro-domain. The WT amino acid residues (D85 and R175), involved in these variations, are highly conserved across the species, suggesting their essential role. Multiple bioinformatic tools, also predicted deleterious nature of these two pro-domain variations. Our *in vitro* functional studies, particularly luciferase reporter assay, demonstrated notable reduction in the activity of downstream promoters such as *Id1*, *Id3*, *Tlx2* and (*SBE)_4_* activities, which is further potentiated to significant level, when co-transfected with *ALK2*. BMP7, like other members of TGF-β superfamily, is synthesized with a large N-terminal pro-domain (254 aa) and a C-terminal mature domain (139 aa). The disulfide-linked mature domain of BMP7 is secreted in a homo/hetero-dimeric form, following intracellular cleavage of pro-domain. However, the BMP7-propeptide remain non-covalently associated with mature dimeric peptide, which is known to stabilize the dimeric ligand and facilitate the extracellular storage, prior to binding to the membrane bound BMP-receptors for further downstream signal (Gregory *et al*., 2005). Accumulating evidences suggested that pro-domain of BMP7 is essential for targeting the dimeric mature domain complex to ECM, where it facilitates the binding of complex (aa Gly74-Arg184) to Fibrillins (Gregory *et al*., 2005; Wohl *et al*., 2016), and it also provide stability to the mature complex. The variant p.D85V localised to α_2_ helix of PD, which is likely involved in mature domain binding and stability. Moreover, PD inhibits type II receptor binding leading to latency of the ligand. Our *in silico* analyses suggest that the variation D85V not only caused perturbation in the 2^nd^ α helix, but also in the extended region of PD (absence of β sheet aa 264-267), thus likely debilitating the protein function. It is also speculated to perturb MMP cleavage site (Furlan *et al*., 2021). Since, beta-strands are considered critical components of the protein’s secondary structure, any alterations in their length or integrity can significantly affect the overall stability and function of the protein. Further, the absence of the beta-strand close to the cleavage site may lead to improper folding or loss of structural rigidity, compromising the protein’s ability to perform its normal functions. The superimposed structure of mutein p.D85V over WT exhibited a RMSD value of 3.016 Å. Similarly, the variant p.R175W that resides in the fibrillin 1 binding site (Gly74 – Arg 184) (Wohl *et al*., 2016), showing conformational changes (as shown in secondary and tertiary structure prediction) likely impair PD dimerization and mature complex formation. The RMSD value of 1.682 Å for superimposed 3D structure for p.R175W, implicate that Tryptophan substitution at 175^th^ position induce large deformities in the tertiary structure of mutein which may affect the folding, liganding and functional modulation of the protein (Boon *et al*., 2020).

The mutein p.A283T lie adjacent to the cleavage site of BMP7. Intracellular cleavage of PD from mature BMP7 peptide by proprotein convertase (PACE) is dependent on certain conserved basic aa residues. Distortion/masking of PACE site may lead to over expression of BMP7 possibly due to defective processing of BMPs (Swencki-Underwood *et al*., 2008). In this study, our *in silico* analysis demonstrate loss of two β-sheets in secondary structure and molecular docking study have shown distorted tertiary structure with high RMSD value (3.501 Å). A competitive displacement of pro-domain and activation of downstream signaling through mature domain-receptor complex has been demonstrated in an elegant study by Sengle et al., 2008 (Sengle *et al*., 2008). *In vitro* gene expression study reveals overexpression of BMP7 as well as BMP2 (a heterodimer forming partner of BMP7), due to p.A283T. Similarly, upregulation of a panel of downstream target genes namely *Nkx2-5, Mef2c, Irx4, Gata4* and co-transcriptional activator *Smad1, Smad5* and *Smad 4* has been noted. Moreover, luciferase assays have shown enhancement of downstream promoter activities in response to A283T variation, neighbouring the cleavage site. However, contrary to our observation, cleavage site mutation p.R316S in BMP9 caused significant decrease in BRE-luc (BMP responsive element) promoter activity and demonstrated reduced mature domain in supernatant of HEK293 cell culture (Wang *et al*., 2019). Similarly, Upton et al. 2022 (Upton *et al*., 2023) described p.S320C mutation causing impaired processing of BMP9 (Upton *et al*., 2023).

Two other mature domain muteins (p.M315I and p.N321S) also cause significant increase in *BMP2/7* expression as well as escalation in the activity of downstream promoters (*Id1, Id3, Tlx2 and SBE_4_*). Moreover, enhancement in the expression (qRT-PCR) of certain other downstream target genes viz., *Gata4*, *Smad1, Smad5* and *Smad4* as well as augmented SMAD1/5 phosphorylation caused due to these C-terminal muteins are also being supported by the previous report by Swencki-Underwood et al., 2008 (Swencki-Underwood *et al*., 2008), where three C-terminal variation cause increase in BMP7 expression 3-10 fold more. We further noted that the expression of *Mef2c, Nkx2-5* and *Irx4* is enhanced in response to p.M315I variant while p.N321S showed a decreasing trend, though certain other abovementioned genes show upregulation. Nonetheless, in case of variants (p.M315I), substitution of methionine (negatively charged) by isoleucine (neutral) affects the overall net charge of BMP7 which might increase stable beta-strand formation and enhanced secretion of mature-domains (Qiao *et al*., 2017); which likely upregulate the downstream signaling.

Further, a remarkable change is marked in the tertiary structure of mutein p.M315I when compared with the WT with RMSD value of 3.568 Å. Conversely, in case of substitution p.N321S, both Asn and Ser are polar, uncharged aa, which do not bring any major conformational change. However, the superimposed structures of p.N321S mutein over WT, resulted a high RMSD value 5.78 Å, further suggesting marked conformational changes. (Hirono *et al*., 2019).

Interestingly, our qRT-PCR data demonstrate down-regulation of some of the LR-patterning genes i.e., *Nodal* **(Figure 5j)***, Pitx2c* **(Figure 5k)** and BMP antagonist, *Chordin* **(Figure 5l)** due to all the muteins compared to WT-BMP7. The antagonistic role of BMP versus Nodal define the L-R axis patterning (Luo and Su, 2012; Lenhart *et al*., 2013) and it is reported by Ramsdell, 2005(Ramsdell *et al*., 2005) that BMP signaling inhibited expression of Nodal in right-lateral plate mesoderm (Ramsdell *et al*., 2005). Our gene expression analysis has shown that the expression of *Nodal* and *Pitx2* are reduced in response to all the five variants due inhibitory effect of BMPs. Further, the expression of *chordin* which is a BMP antagonist, has also observed to be decreased due to all the five variants.

In the present study, the expression of Bmp7 (Western blot) is depicted to show no significant change in P19 cell extracts, transfected with either WT-BMP7 or due to any of five variants (D85V, R175W, A283T, M315I and N321S). However, all these five mutants caused increased expression of many downstream target genes. It is well established that BMP7 form heterodimer either with BMP2 or BMP4 (Suzuki *et al*., 1997) which is more potent than their respective homodimer. Moreover, some studies reported that the biological activity of heterogenous dimers is almost 20 times higher than that of homodimers (Aluganti Narasimhulu and Singla, 2020). Furthermore, studies in Xenopus and Zebrafish has also revealed the enhanced activity of these heterodimers in embryonic assays (Suzuki *et al*., 1997; Schmid *et al*., 2000). In mammalian system (mice model), BMP7 predominantly function as heterodimer with either BMP2 or BMP4, which has been confirmed by co-immunoprecipitation assay (Kim *et al*., 2019). In light of this, when the expression of *Bmp7* and *Bmp2* was checked, elevated levels of expression of both the genes are detected in response to all the five variations compared to WT. Since functional redundancy of these proteins is well known, co-operative heterodimer formation might result in increased activity of downstream target genes. This can be correlated with GoF mutation reported in ALK2 (Groppe *et al*., 2023). Hence, it is presumed that the overexpression of both the heteromeric BMPs, likely circumvent the BMP-receptor mediated downstream signaling, thus up-regulating many of the target genes as seen in our case.

Extensive survey of literature, indicate that BMP7 influence cell proliferation and migration in a context dependent manner (Alarmo *et al*., 2009). BMP7 potentially repair damaged renal epithelial cells (Zeisberg *et al*., 2003). Further, BMP signaling play crucial role during cardiomyocyte proliferation and heart development (Yuasa and Fukuda, 2008; Wang *et al*., 2014). In order to evaluate the effect of BMP7 muteins on cell proliferation we checked the mitotic index of H9c2 cells transfected with WT and mutants of BMP7 and observed a significant increase in mitotic index in all the five muteins, compared to WT and this could be correlated with the GoF activity of the muteins in all other aforementioned assays. In a mouse model study (Kim *et al*., 2001) have shown that Bmp6;Bmp7 double mutant had high mitotic index in myocardium with atrial and ventricular septal defects due to abnormal growth regulation (Kim *et al*., 2001). Further, Bongiovanni et al., 2024 (Bongiovanni *et al*., 2024) also revealed that *Bmp7* GoF increases cardiomyocyte proliferation in zebra fish (Bongiovanni *et al*., 2024).

Further, the down regulation of L/R patterning genes (*Nodal* and *Pitx2*) and the BMP antagonist (*Chordin*) also revealed that these mutants are overexpressing. Interestingly, enhanced trafficking and cell proliferation also supporting that these mutants are working like GoF mutations.

It is noteworthy that gain of BMP function in Noggin^-/-^ (a dedicated BMP antagonist) and Smad6^-/-^ (Bmp-specific nuclear inhibitor) mice similarly result in thickened or hyperplastic valve (Conway *et al*., 2011). Further, GoF mutation in *TGFβ1* (Yadav *et al*., 2022) and *CITED2* (Yadav *et al*., 2021) has also been associated with CHD. On wholesome, we could infer that heterogenous phenotypes of CHD observed due to mutations in different domains of *BMP7* and GoF mutations affecting the BMP signaling pathway during different developmental stages of cardiogenesis.

## 5. Conclusion

In brief, here we identified five non-synonymous rare variations in BMP7, in a large cohort of non-syndromic CHD cases with diverged cardiac phenotype. To the best of our knowledge, this is the first case-control study showing association of *BMP7* with isolated CHD phenotype. *In vitro* and *in silico* characterization all the variations, whether belonging to pro-domain or mature domain not only depicted unique GoF activity of BMP signaling when mutated, but also highlighted its critical role during heart development. Conway et al, 2011 previously hypothesised that either loss-of/ gain-of-function in BMP might be associated with cardiac defects (Conway *et al*., 2011). BMP signaling is also known to have crosstalk with other signalings such as NODAL, TGF-β (Kishigami and Mishina, 2005; Chocron *et al*., 2007) NOTCH, WNT/β-CATENIN (Lin and Hankenson, 2011) pathways during EMT which clearly suggest that there is complex interaction/counteraction among multiple members of signaling network which culminate in to clinically varied CHD phenotype. Besides, the influence of environmental factors could not be ruled out. Alcohol exposure is also shown to boosts the BMP signaling by enhancing the H3 acetylation of cardiac enrich genes *NKX2.5, GATA4*, and *MEF2C* which is presumed to induce CHD (Shi *et al*., 2017). Provided the current evidences of association of *BMP7* mutations with isolated CHD, further studies on gene-gene interaction and gene-environment interaction, within the signaling network would likely shed light on mechanistic insight into cardiac developmental process.

## Acknowledgment

We are very thankful to all the patients, their family members and control individuals for their participation in the present study. We are extremely grateful to Prof. T.K. Lahiri and Dr. Damyanti Agrwal from Department of Cardio-vascular and Thoracic Surgery, IMS, BHU, Varanasi for their constant support and encouragement in enrolment of patients. We would like to acknowledge Dr. Ramkumar Sambasivan, InStem, Bengaluru, Karnataka, India for providing P19 cells, Dr. Maria Genander, Karolinska Institutet, Stockholm, Sweden for *Id1* luciferase reporter construct, Prof. Daniel J. Bernard, Quebec, Canada for *Id3* luciferase reporter construct, Dr. Peter ten Dijke, Netherlands for p(SBE)4-luc reporter plasmid and Prof. Jeffrey L. Wrana, University of Toronto, Canada for Tlx2 reporter construct. We would also like to thanks Dr. Anand Kumar Singh, Department of Zoology, IISER Tirupati for his valuable assistance and guidance in Confocal Microscopy. We are very grateful to Prof. Rajiva Raman, Department of Zoology, BHU, Varanasi, India for critically reviewing the manuscript, his valuable suggestions and grammatical corrections. Thanks are also extended to University Grants Commission (UGC) for junior and senior research fellowship (577/(CSIR-UGC NET JUNE 2018) to Jyoti Maddhesiya. Indian Council of Medical Research (ICMR), Ministry of Health, Govt. of India, New Delhi was also thanked for senior research fellowship (45/4/2013/CMB-BMS) to Dr. Ritu Dixit.

## Conflict of Interest

The authors declare no conflict of interest. All authors have read the manuscript and approved the submission of current version of the manuscript.

## Data Availability

Raw data and derived data supporting the findings of this study are available from the corresponding author on request.

## CRediT authorship contribution

**Bhagyalaxmi Mohapatra:** Conceptualization, Data curation, Formal analysis, Funding acquisition, Investigation, Project administration, Resources, Supervision, Visualization, Writing – review & editing.

**Jyoti Maddhesiya**: Conceptualization, Data curation, Formal analysis, Methodology, Software, Validation, Visualization, Writing – original draft, Writing – review & editing.

**Ritu Dixit:** Conceptualization, Data curation, Formal analysis, Methodology, Visualization, Writing – review & editing.

**Ashoka Kumar:** Investigation

## Source of funding

This study was partially sponsored by Department of Biotechnology (DBT), Govt. of India (grant number-BT/PR14501/MED/12/479/2010) and Institute of Eminence (IOE) fund to BHU by Govt. of India. The funding agency had no involvement in the study design, sample collection, data analysis or interpretation, manuscript preparation, or the decision to submit the article for publication.

## Notes

### Competing Interest Statement

The authors have declared no competing interest.

