## Supplementary material for "Deciphering the functional association of novel variants of *BMP7* in isolated congenital heart disease by integrating *in vitro* and *in silico* approaches": Supplemeatary Material

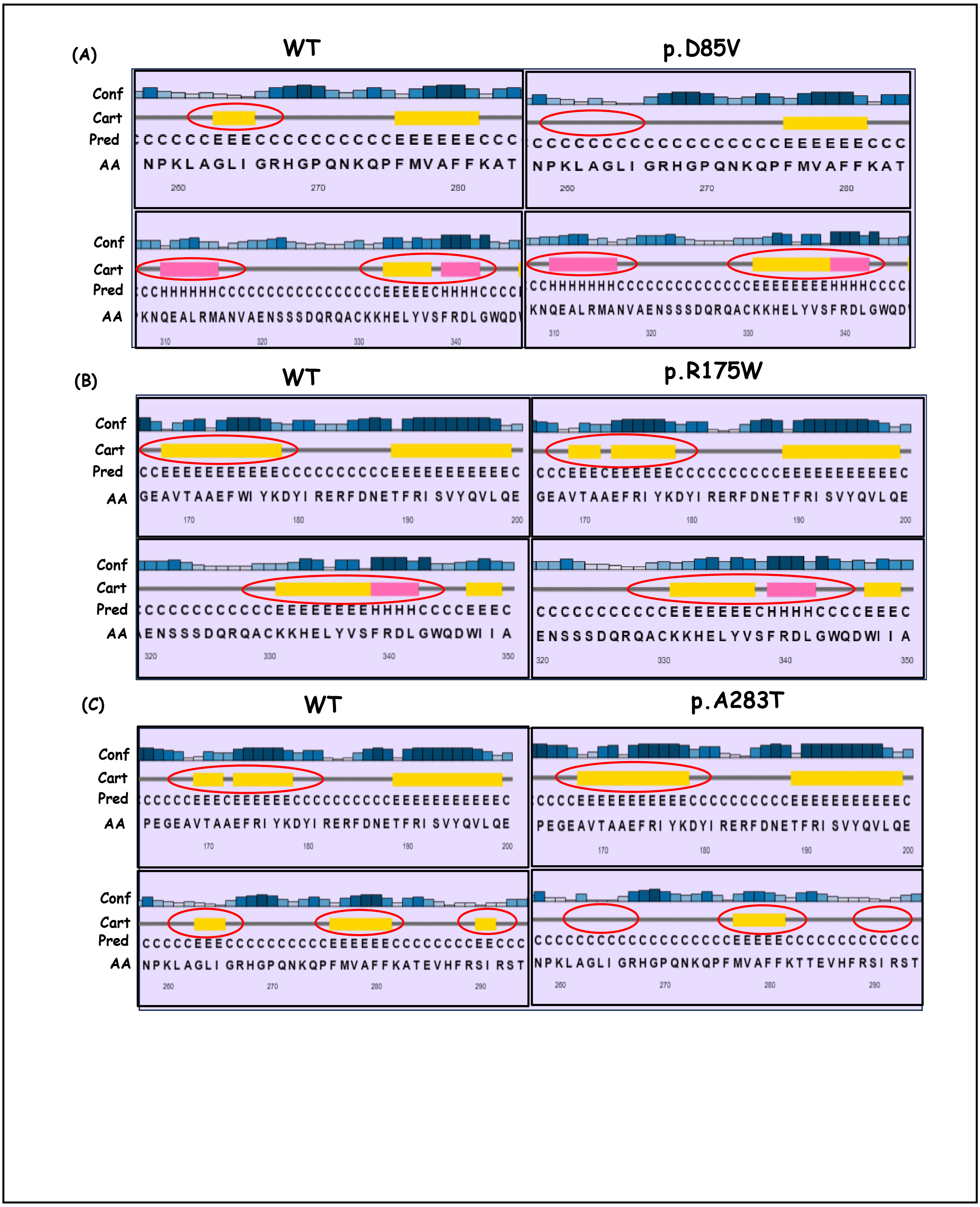
(A)
WT
p.D85V
(B)
WT
p.R175W
(C)
WT
p.A283T
Conf
Cart
Pred
AA

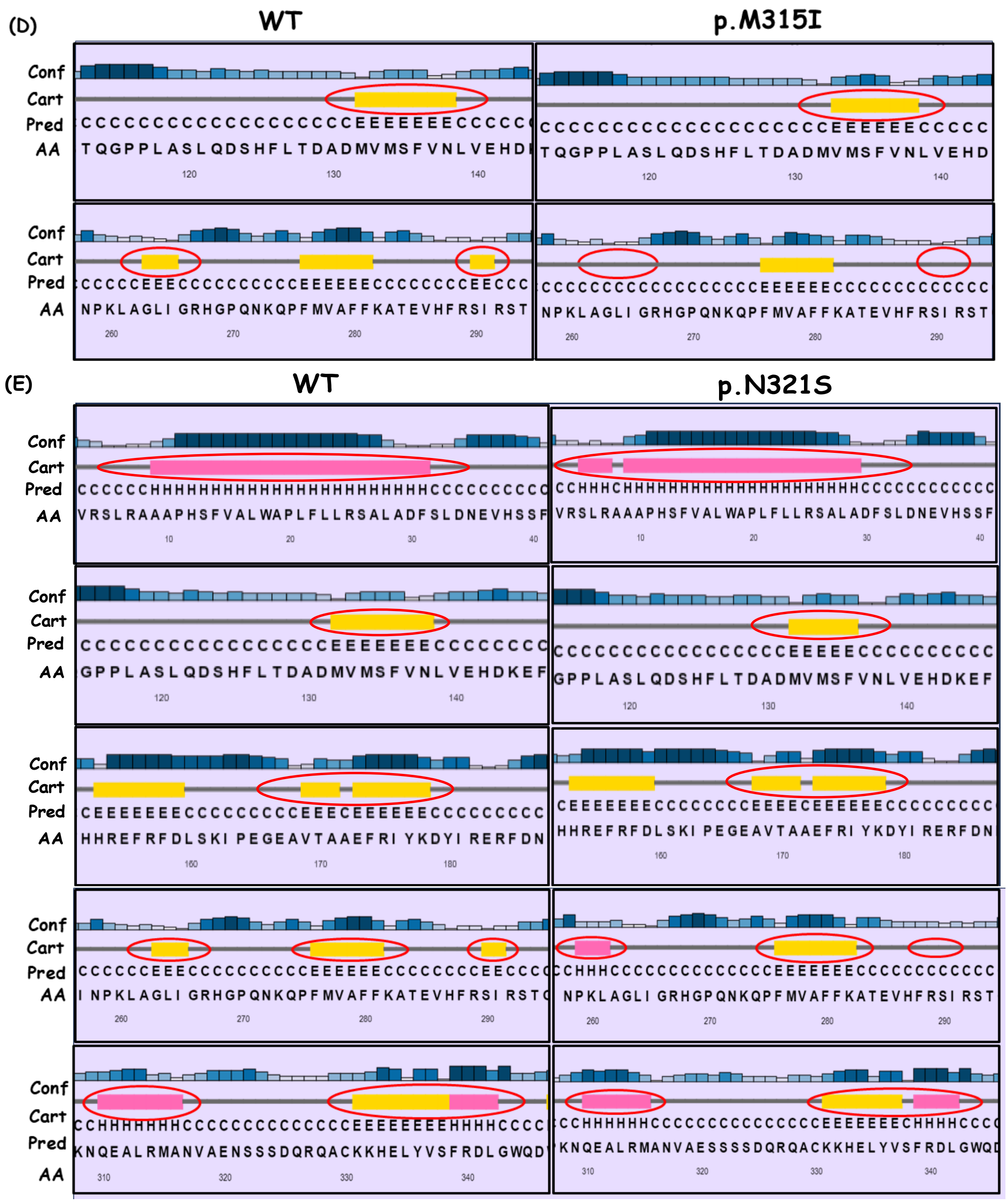


**Supplementary Figure1.** Comparative analysis of secondary structures of BMP7 wild-type (WT) and muteins (MUT) by Psipred server. (A-E) Predicted secondary structures of BMP7 wild-type and muteins (p.D85V, p.R175W, p.A283T, p.M315I and p.N321S). The alterations in structures- helix (pink boxes) and strand (yellow boxes) has been encircled in both WT and MUTs.

**Supplementary Table S1.** Clinical phenotype with their relative frequency in male and females

| CHD | CHD Phenotype | Number (%) |
|---|---|---|
| **Acyanotic defects** | Ventricular septal defects (VSD) | 93 (32.6) |
| | Atrial septal defects (ASD) | 59 (20.7) |
| | ASD, VSD | 18 (6.3) |
| | Patent ductus arteriosus | 7 (2.4) |
| | Atrio-ventricular canal defect | 6 (2.1) |
| | Eisenmenger syndrome | 4 (1.4) |
| | Gerbode defect | 2 (0.7) |
| | Patent foramen ovale | 1 (0.3) |
| **Cyanotic defects** | Tetralogy/ Pentalogy of Fallot | 44 (15.4) |
| | Transposition of great arteries | 6 (2.1) |
| | Dextrocardia | 6 (2.1) |
| | Tricuspid Atresia | 3 (1.05) |
| | Ebstein Anomaly | 3 (1.05) |
| | Double outlet right ventricle | 2 (.07) |
| | Total/Partial Pulmonary Venous Connection | 2 (.07) |
| **Left/ right obstructive defects** | Bicuspid aortic valve | 12 (4.2) |
| | Pulmonary stenosis | 6 (2.1) |
| | Aortic stenosis | 3 (1.05) |
| | Truncus Arteriosus | 3 (1.05) |
| | Coarctation of aorta | 1 (0.3) |
| **Others** | Holt Oram Syndrome | 2 (0.7) |
| | Single Left ventricle | 1 (0.3) |
| | Double aortic arc | 1 (0.3) |
| **Total** | | **285** |

**Supplementary Table S2**. Position of changed α-helix and β-sheet caused due to identified variants of *BMP7* as predicted by Psipred server

| Variants | α-helix | β-sheet |
|---|---|---|
| p.D85V | 310-316th extension | 331-338th extension<br>264-267th lost |
| p.R175W | - | 168-178th shortening<br>331-338th shortening |
| p.A283T | - | 263-265th lost<br>290-291st lost<br>169-171th extension |
| p.M315I | - | 132-138th shortening<br>263-265th lost<br>290-291st lost |
| p.N321S | 9-31st shortening<br>259-261st addition<br>310-316th shortening | 132-138th shortening<br>168-178th extension<br>263-265th lost<br>276-281st extension<br>331-338th shortening<br>264-265th addition<br>148-150th shortening |
